# Phylogenomics of a new fungal phylum reveals multiple waves of reductive evolution across Holomycota

**DOI:** 10.1101/2020.11.19.389700

**Authors:** Luis Javier Galindo, Purificación López-García, Guifré Torruella, Sergey Karpov, David Moreira

## Abstract

Compared to multicellular fungi and unicellular yeasts, unicellular fungi with free-living flagellated stages (zoospores) remain poorly known and their phylogenetic position is often unresolved. Recently, rRNA gene phylogenetic analyses of two atypical parasitic fungi with amoeboid zoospores and long kinetosomes, the sanchytrids *Amoeboradix gromovi* and *Sanchytrium tribonematis*, showed that they formed a monophyletic group without close affinity with known fungal clades. Here, we sequence single-cell genomes for both species to assess their phylogenetic position and evolution. Phylogenomic analyses using different protein datasets and a comprehensive taxon sampling result in an almost fully-resolved fungal tree, with Chytridiomycota as sister to all other fungi, and sanchytrids forming a well-supported, fast-evolving clade sister to Blastocladiomycota. Comparative genomic analyses across fungi and their allies (Holomycota) reveal an atypically reduced metabolic repertoire for sanchytrids. We infer three main independent flagellum losses from the distribution of over 60 flagellum-specific proteins across Holomycota. Based on sanchytrids’ phylogenetic position and unique traits, we propose the designation of a novel phylum, Sanchytriomycota. In addition, our results indicate that most of the hyphal morphogenesis gene repertoire of multicellular fungi had already evolved in early holomycotan lineages.

## Introduction

Within the eukaryotic supergroup Opisthokonta, multicellularity evolved independently in Fungi and Metazoa from unicellular ancestors along their respective branches, Holomycota and Holozoa^1^. The ancestor of this major supergroup originated ~1-1,5 Ga ago^2–4^ and likely possessed one posterior flagellum for propulsion in aquatic environments^1^. This character has been retained in many modern fungal lineages at least during some life cycle stages^5,6^. Along the holomycotan branch, the free-living, non-flagellated nucleariid amoebae were the first to diverge, followed by the flagellated, phagotrophic, endoparasitic Rozellida (Cryptomycota)^7–9^ and Aphelida^10^, and the highly reduced, non-flagellated Microsporidia^11,12^. Aphelids branch as sister lineage to *bona fide*, osmotrophic, Fungi^13^ (i.e., the zoosporic Blastocladiomycota and Chytridiomycota, and the non-zoosporic Zoopagomycota, Mucoromycota, Glomeromycota, and Dikarya). Within fungi, except for the secondary flagellar loss in the chytrid *Hyaloraphydium curvatum* ^14^, all known early divergent taxa are zoosporic (having at least one flagellated stage).

Zoosporic fungi are widespread across ecosystems, from soils to marine and freshwater systems, from tropical to Artic regions^15,16^. They include highly diverse saprotrophs and/or parasites, participating in nutrient recycling through the “mycoloop”^17–20^. Initially considered monophyletic, zoosporic fungi were recently classified into Blastocladiomycota and Chytridiomycota^21^, in agreement with multigene molecular phylogenies^13,21,22^. These two lineages appear sister to the three main groups of non-flagellated fungi, Zoopagomycota, Mucoromycota and Dikarya, for which a single ancestral loss of the flagellum has been proposed^23^. Characterizing the yet poorly-known zoosporic fungi is important to understand the evolutionary changes (e.g., flagellum loss, hyphae development) that mediated land colonization and the adaptation of fungi to plant-dominated terrestrial ecosystems^24,25^. This requires a well-resolved phylogeny of fungi including all zoosporic lineages. Unfortunately, previous phylogenomic analyses did not resolve which zoosporic group, either Blastocladiomycota or Chytridiomycota, is sister to non-flagellated fungi^21,26–28^. This lack of resolution may result from the old age of these splits (~0,5-1 Ga)^3,29,30^ and the existence of several radiations of fungal groups, notably during their co-colonization of land with plants^25,31^, which would leave limited phylogenetic signal to resolve these deep nodes^22^.

Because of this phylogenetic uncertainty, the number and timing of flagellum losses in fungi remain under debate, with estimates ranging between four and six for the whole Holomycota^32^. Improving taxon sampling with new, divergent zoosporic fungi can help resolving these deep nodes. One of such lineages is the sanchytrids, a group of chytrid-like parasites of algae with uncertain phylogenetic position represented by the genera *Amoeboradix* and *Sanchytrium*, which exhibit a highly reduced flagellum with an extremely long kinetosome^33,34^ (Fig. 1). Determining the phylogenetic position of this zoosporic lineage remains decisive to infer the history of flagellum losses and the transition to hyphal-based multicellularity.

**Fig. 1.**
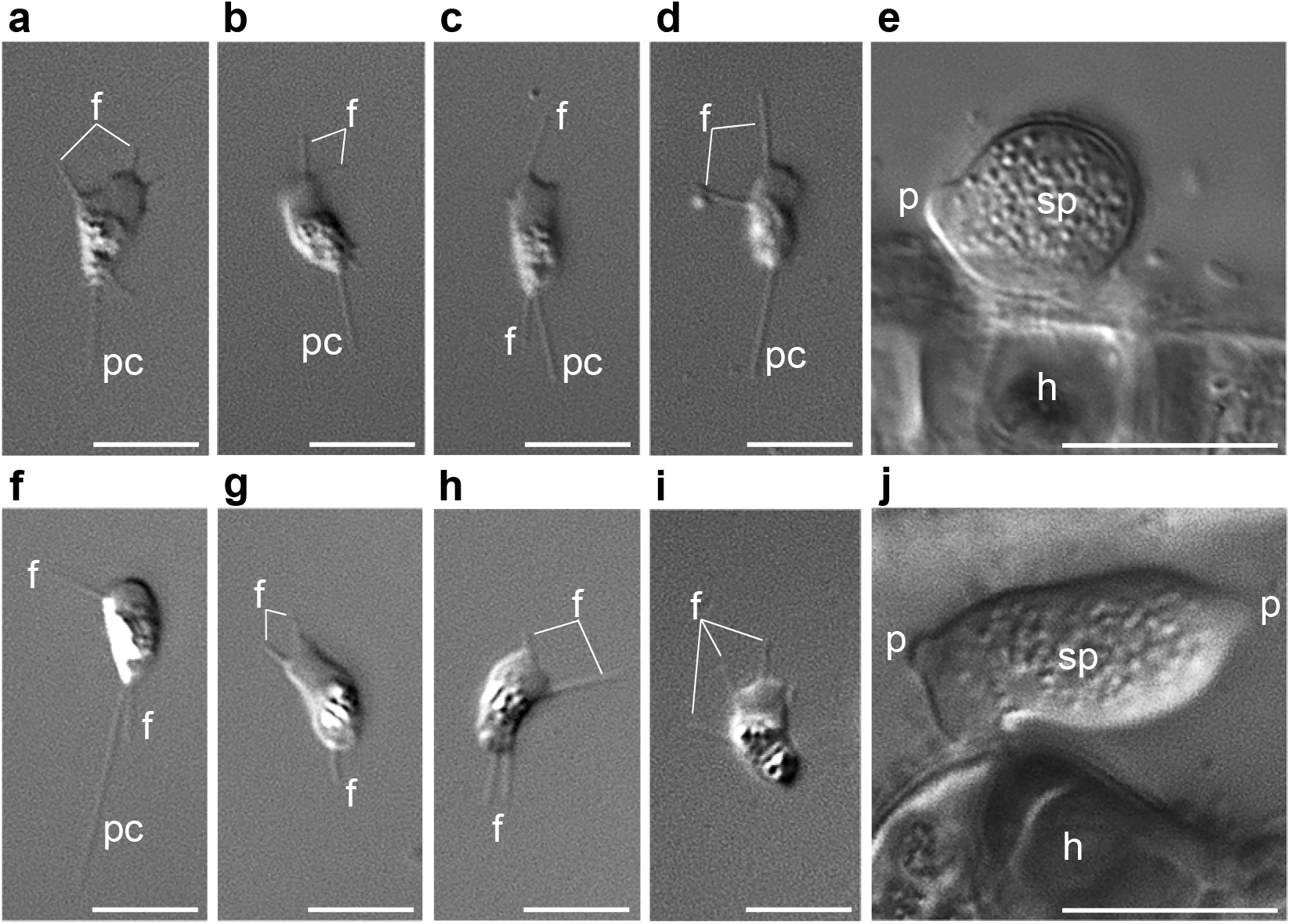
Light microscopy images of life cycle stages of *Sanchytrium tribonematis* (a-e) and *Amoeboradix gromovi* (f-j). **a-d**, **f-i** – amoeboid crawling zoospores with filopodia (f) and posterior pseudocilium (pc). **g-i** – zoospores with retracted pseudocilium. **e, j** – sporangium (sp) with one (**e**) or two (**j**) papillae (p) on the host (h) surface. Scale bars: **a-d**, **f-i** – 5 μm; **e, j** – 10 μm.

In this work we generate the first genome sequences for the sanchytrids *Amoeboradix gromovi* and *Sanchytrium tribonematis* and analyse them together with available genomic and transcriptomic data for Chytridiomycota and Blastocladiomycota. We obtain an almost fully-resolved phylogeny of fungi which shows sanchytrids as a new fast-evolving lineage sister to Blastocladiomycota. Contrasting with previous weakly-supported analyses^13,22,28,35,36^, we robustly place the root of the fungal tree between chytrids and all other fungi. Our new phylogenomic framework of Fungi supports a conservative model of three flagellum losses in Holomycota and highlights the importance of early-diverging unicellular Holomycota in the evolution of hyphal-based multicellularity.

## Results and discussion

### The new zoosporic fungal phylum Sanchytriomycota

We isolated individual sporangia of *Amoeboradix gromovi* and *Sanchytrium tribonematis* (Fig. 1) by micromanipulation and sequenced their genomes after whole genome amplification. After thorough data curation (see Methods), we assembled two high-coverage genome sequences (123.9X and 45.9X, respectively) of 10.5 and 11.2 Mbp, encoding 7,220 and 9,638 proteins, respectively (Table 1). Comparison with a fungal dataset of 290 near-universal single-copy orthologs^37^ indicated very high completeness for the two genomes (92.41% for *A. gromovi*; 91.72% for *S. tribonematis*). Only half of the predicted sanchytrid proteins could be functionally annotated using EggNOG^38^ (3,838 for *A. gromovi*; 4,772 for *S. tribonematis*) (Supplementary Data 1). This could be partly due to the fact that, being fast-evolving parasites^33,34,39^, many genes have evolved beyond recognition by annotation programs^40,41^. However, low annotation proportions are common in Holomycota, including fast-evolving parasites (e.g., only 20% and 52% of the genes of the microsporidia *Nosema parisii* and *Encephalitozoon cuniculi* could be assigned to Pfam domains and GO terms^40,42^) but also the less fast-evolving metchnikovellids (*Amphiamblys* sp., 45.6%), rozellids (*Rozella allomycis*, 64.9%; *Paramicrosporidium saccamoebae*, 66.7%) and blastocladiomycetes (*Catenaria anguillulae*, 47.5%). Many of the non-annotated genes were unique to sanchytrids as deduced from orthologous gene comparison with 57 other species, including representatives of the other major fungal lineages, and several outgroups (Supplementary Data 2). After clustering of orthologous proteins with OrthoFinder ^43^, we identified 1217 that were only present in both *A. gromovi* and *S. tribonematis*. Their analysis using eggNOG^38^ resulted in only 93 proteins annotated. The remaining (93.4%) sanchytrid-specific proteins lack functional annotation (Supplementary Data 2).

**Table 1.** Statistics of sanchytrid genome assemblies before and after decontamination and comparison with related zoosporic lineages.

|  | *Amoeboradix gromovi* |  | *Sanchytrium tribonematis* |  | *Allomyces macrogynus* | *Catenaria anguillulae* | *Rhizoclosmatium globosum* | *Spizellomyces punctatus* | *Anaeromyces robustus* |
| --- | --- | --- | --- | --- | --- | --- | --- | --- | --- |
|  | Before | After | Before | After |  |  |  |  |  |
| Genome size (Mb) | 48.7 | 10.5 | 37.1 | 11.2 | 52.62 | 41.34 | 57.02 | 24.1 | 71.69 |
| GC% | 51.06 | 36.27 | 42.9 | 34.64 | 61.6 | 56 | 44.9 | 47.6 | 16.3 |
| Number of contigs | 8,420 | 1,167 | 8,015 | 1,960 | 8,973 | 509 | 437 | 329 | 1,035 |
| N50 | 27,236 | 13,376 | 17,350 | 11,874 | 35,497 | 217,825 | 292,246 | 155,888 | 141,798 |
| Predicted proteins | 87,868 | 7,220 | 50,780 | 9,368 | 19,447 | 12,763 | 16,987 | 9,422 | 12,083 |
| Mt genome size (bp) | 27,055 |  | 24,749 |  | 57,473 | --- | --- | 126,474 | --- |
| Mt genome GC% | 30.69 |  | 25.86 |  | 39.5 | --- | --- | 29.88 | --- |

The two sanchytrid genomes yielded similar sequence statistics (Table 1) but showed important differences with genomes from other well-known zoosporic fungi. They are 4-5 times smaller than those of blastocladiomycetes (40-50 Mb) and average chytrids (~20 to 101 Mb), an observation that extends to the number of protein-coding genes. Their genome G+C content (~35%) is much lower than that of blastocladiomycetes and most chytrids (40-57%, though some chytrids, like *Anaeromyces robustus*, may have values down to 16.3%^44,45^). Low G+C content correlates with parasitic lifestyle in many eukaryotes^46^. In Holomycota, low G+C is observed in microsporidian parasites and Neocallimastigomycota, both anaerobic and exhibiting reduced mitochondrion-derived organelles^45,47^, and in the aerobic parasite *R. allomycis*^9^. Although sanchytrids are aerobic parasites with similar life cycles to those of blastocladiomycetes and chytrids^33,34,39^, their smaller genome size and G+C content suggest that they are more derived parasites. This pattern is accompanied by a global acceleration of evolutionary rate (see below), a trend also observed, albeit at lower extent, in *R. allomycis*^9,48–50^.

The mitochondrial genomes of *S. tribonematis* and *A. gromovi* showed similar trends. Gene order was highly variable (Supplementary Fig. 1), as commonly observed in Fungi^51^, and their size (24,749 and 27,055 bp, respectively) and G+C content (25.86% and 30.69%, respectively) were substantially smaller than those of most other Fungi. However, despite these signs of reductive evolution, most of the typical core mitochondrial genes were present, indicating that they have functional mitochondria endowed with complete electron transport chains.

To resolve the previously reported unstable phylogenetic position of sanchytrids based on rRNA genes^33^, we carried out phylogenomic analyses on a manually curated dataset of 264 conserved proteins (93,743 amino acid positions)^13,49,52^ using Bayesian inference (BI) and maximum likelihood (ML) with the CAT^53^ and PMSF^54^ models of sequence evolution, respectively. Both mixture models are known to alleviate homoplasy and long-branch attraction (LBA) artefacts^22,53^. We used the same 59 species as above (dataset GBE59), including a wide representation of holomycota plus two holozoa, two amoebae and one apusomonad as outgroups. BI and ML phylogenomic analyses yielded the same tree topology for major fungal groups with only minor changes in the position of terminal branches (Fig. 2a; Supplementary Fig. 2a-b). We recovered maximum statistical support for both the monophyly of sanchytrids (*A. gromovi* + *S. tribonematis*) and their sister position to Blastocladiomycota. Thus, sanchytrids form a new deep-branching zoosporic fungal clade. Given their divergence and marked genomic differences with their closest relatives (Blastocladiomycota), we propose to create the new phylum Sanchytriomycota to accommodate these fungal species (see description below).

**Fig. 2.**
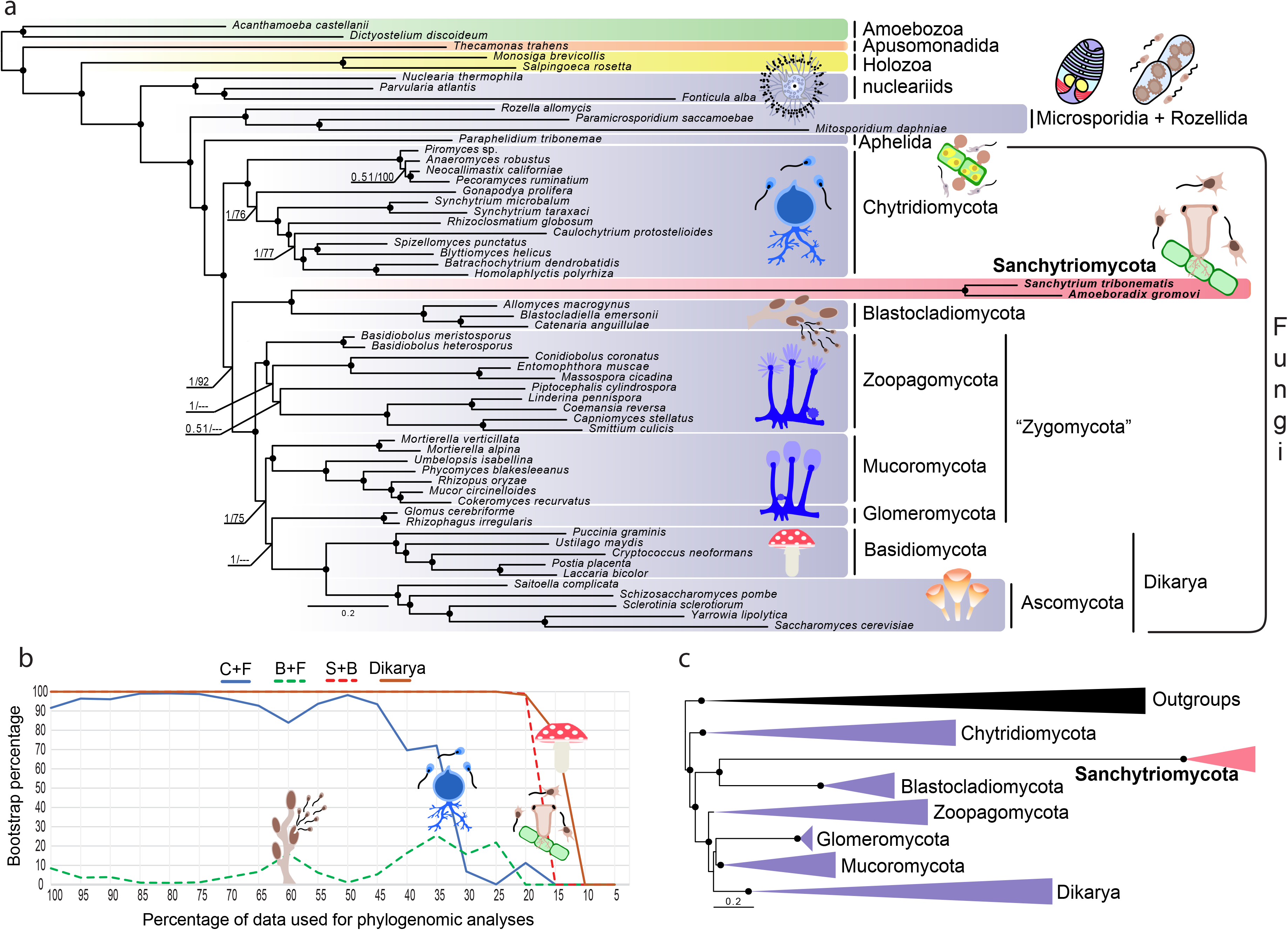
Phylogenomic analysis of Holomycota. **a**, Bayesian inference (BI) phylogenomic tree based on the GBE dataset of 264 conserved proteins (93,743 amino acid positions). The tree was reconstructed using 59 species and the CAT-Poisson model and the PMSF approximation of the LG+R8+C20 model for maximum likelihood (ML). **b**, Evolution of IQ-TREE ML bootstrap support for Chytridiomycota sister to all other fungi (C+F), Blastocladiomycota+Sanchytriomycota sister to all other fungi (B+F), Sanchytriomycota within Blastocladiomycota (S+B), and the monophyly of Dikarya (Dikarya) as a function of the proportion of fast-evolving sites removed from the GBE59 dataset. Holomycota are highlighted in violet, sanchytrids in pink and outgroup taxa in other colors (green, orange and yellow). All phylogenomic trees can be seen in Supplementary Figs. 2a-e. **c**, Schematic BI phylogeny showing the results obtained with the BMC dataset of 53 conserved proteins (14,965 amino acid positions) using 59 species (Fungi: violet; Sanchytriomycota: red; outgroup taxa: black) and the CAT-GTR model (BI) and the PMSF approximation of the LG+R7+C60 model for maximum likelihood (ML). In both trees, branches with support values higher or equal to 0.99 BI posterior probability and 99% ML bootstrap are indicated by black dots.

### An updated phylogeny of Fungi

In addition to the monophyly of Sanchytriomycota and Blastocladiomycota, our analysis retrieved chytrids as sister group to all other fungi with full Bayesian posterior probability (PP=1) and smaller ML bootstrap support (BS=92%). However, this position of the root of fungi on the chytrid branch became fully supported (BS=100%) after removing the fast-evolving sanchytrids in a new dataset of 57 species and 93,421 amino acid positions (dataset GBE57; Supplementary Fig. 2c), suggesting an LBA artefact on the tree containing the sanchytrids. In fact, as observed in rRNA gene-based phylogenies^33^, sanchytrids exhibited a very long branch (Fig. 2a), indicating a fast evolutionary rate. Long branches associated with fast-evolving genomes, well-known in Microsporidia^48^ and other Holomycota, can induce LBA artefacts in phylogenetic analyses^55–57^. To further study if LBA affected the position of the fast-evolving sanchytrids and other fungi in our tree, we carried out several tests. First, we introduced in our dataset additional long-branched taxa, metchnikovellids and core Microsporidia, for a total of 74 species and 86,313 conserved amino acid positions (dataset GBE74). Despite the inclusion of these LBA-prone fast-evolving taxa, we still recovered the monophyly of sanchytrids and Blastocladiomycota (Supplementary Fig. 2d). However, they were dragged to the base of Fungi, although with low support (BS 78%). To confirm the possible impact of LBA on this topology, we removed the fast-evolving sanchytrids (dataset GBE72; 72 species and 84,949 amino acid positions), which led to chytrids recovering their position as sister group of all other fungi with higher support (BS 91%; Supplementary Fig. 2e), corroborating the LBA induced by the long sanchytrid branch.

Second, we tested the influence of fast-evolving sites by applying a slow-fast approach^58^ that progressively removed the fastest-evolving sites (in 5% steps) in the 59-species alignment. The monophyly of sanchytrids and Blastocladiomycota obtained maximum support (BS >99%) in all steps until only 20% of the sites remained, when the phylogenetic signal was too low to resolve any deep-level relationship (Fig. 2b). This relationship was as strongly supported as a well-accepted relationship, the monophyly of Dikarya. Similarly, the root of the fungal tree between chytrids and the rest of fungi was supported (>90% bootstrap) until only 40% of the sites remained. By contrast, a root between sanchytrids+Blastocladiomycota and the rest of fungi always received very weak support (<26% bootstrap).

We then further tested the robustness of the position of chytrids using alternative topology (AU) tests. For the dataset GBE59, these tests did not reject alternative positions for the divergence of Chytridiomycota and Blastocladiomycota+Sanchytriomycota (*p*-values >0.05; Supplementary Data 3), which likely reflected the LBA due the long sanchytrid branch. In fact, the position of Blastocladiomycota at the base of fungi was significantly rejected (*p*-values <0.05; Supplementary Data 3) after removing the fast-evolving sanchytrid clade (dataset GBE57).

Finally, we compared the results based on the GBE dataset with those based on a different dataset. We decided to use the BMC dataset^59^ (which includes 53 highly conserved proteins and 14,965 amino acid positions) because it was originally designed to study fast-evolving holomycota (e.g. Microsporidia), which made it appropriate to deal with the fast-evolving sanchytrids. Using the same taxon sampling of 59 species (BMC59), we recovered the same topology, particularly for the deeper nodes, with full ML ultrafast and conventional bootstrap (BS=100) and Bayesian posterior probability (PP=1) supports for both the monophyly of sanchytrids+blastocladiomycetes and chytrids as the sister lineage to all other fungi (Fig. 2c; Supplementary Fig. 2f-h).

The origin of the conspicuous long branch exhibited by sanchytrids is unclear. It has been shown that fast-evolving organisms, including those within Holomycota, tend to lack part of the machinery involved in genome maintenance and DNA repair^50,60^. To verify if it was also the case in sanchytrids, we searched in the two sanchytrid genomes 47 proteins involved in genome maintenance and DNA repair that have been observed to be missing in several fast-evolving budding yeasts^60^. Our results confirm that most of these genes are also absent in both sanchytrids (Supplementary Fig. 3). However, they are also missing in the closely related short-branching blastocladiomycete *A. macrogynus*, suggesting that the long sanchytrid branch is not only due to the absence of these genes.

The relative position of Chytridiomycota or Blastocladiomycota as first branch to diverge within Fungi has remained a major unresolved question^61^. If the earliest fungal split occurred ~1 billion years ago^30^, the phylogenetic signal to infer it may have been largely eroded over time. Likewise, if evolutionary radiations characterized early fungal evolution^22^, the accumulation of sequence substitutions during early diversification would have been limited. Both factors would explain the difficulty to resolve the deepest branches of the fungal tree so far. Our results, based on an improved gene and taxon sampling, provided strong support for the placement of chytrids as sister clade to all other Fungi. This solid position of the root on the chytrid branch is additionally consistent with the distribution of so-considered derived characters in Blastocladiomycota, including sporic meiosis, relatively small numbers of carbohydrate metabolism genes and, in some species, hyphal-like apical growing structures (*Allomyces*) and narrow sporangia exit tubes (e.g., *Catenaria* spp.)^5,62–64^. Despite the use of a large dataset, some branches remained unresolved, in particular the position of Glomeromycota, sister either to Mucoromycota or Dikarya (Fig. 2a), which has important implications related to their symbiotic adaptation to land plants^24,65,66^.

### Macroevolutionary trends in primary metabolism

To assess if sanchytrid metabolic capabilities are as reduced as suggested by their small genome sizes, we inferred their metabolic potential in comparison with other major fungal clades as well as other opisthokonts and amoebozoa as outgroup (43 species). We compared the EggNOG^38^-annotated metabolic repertoires of holomycotan phyla by focusing on 1,158 orthologous groups (Supplementary Data 4) distributed in eight primary metabolism categories. Unexpectedly, cluster analyses based on the presence/absence of these genes did not group sanchytrids with canonical fungi and their closest aphelid relatives (i.e. *Paraphelidium*), but with non-fungal parasites (*R. allomycis, Mitosporidium daphniae* and *P. saccamoebae*) which show evidence of reductive genome evolution^9,11,67^ (Fig. 3a), suggesting gene loss-related convergence. An even further metabolic reduction was observed in Neocallimastigomycota, gut-inhabiting symbiotic anaerobic chytrids^68–70^. A principal coordinate analysis of the same gene matrix confirmed this result (Fig. 3b).

**Fig. 3.**
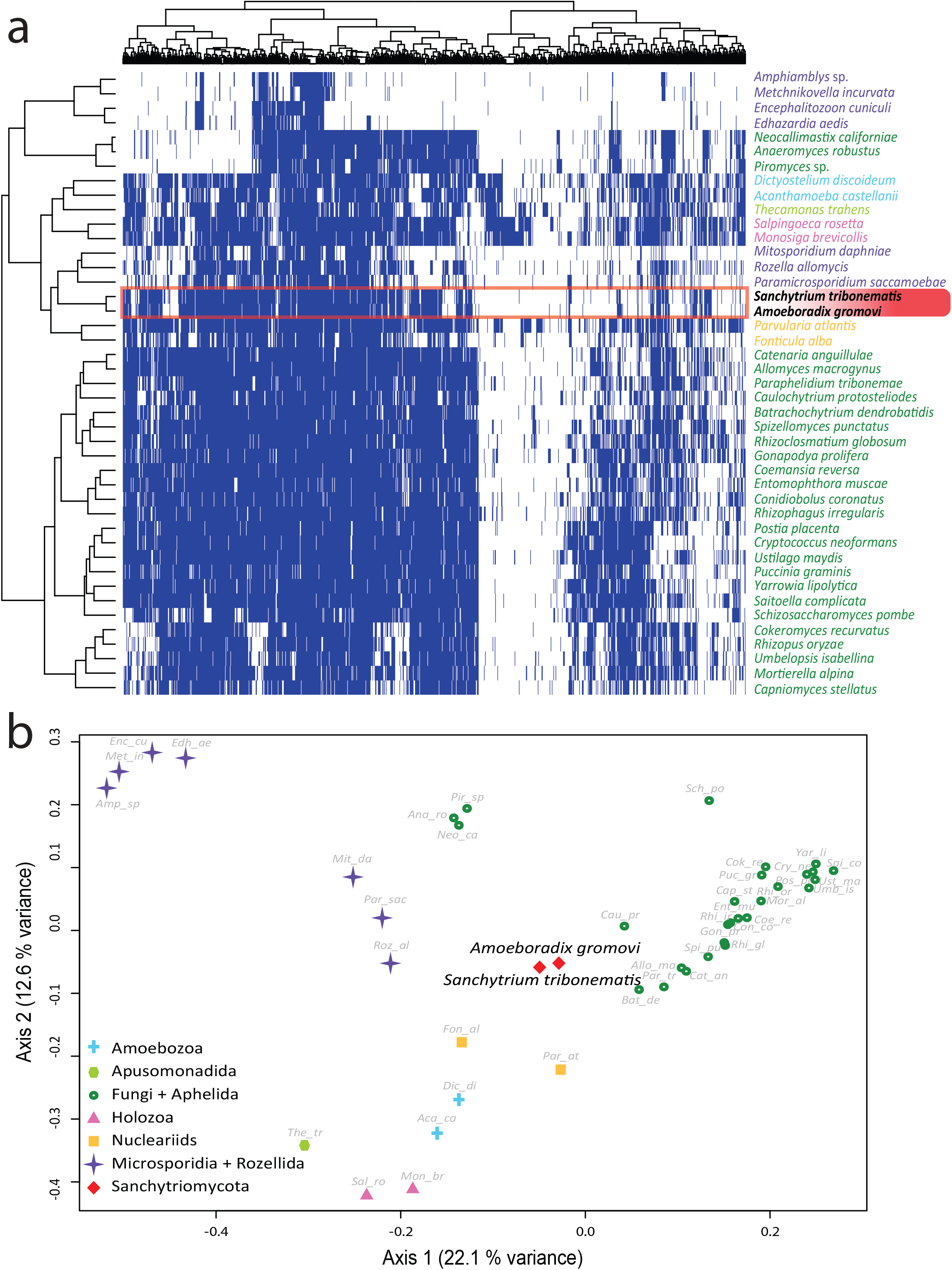
Distribution patterns of primary metabolism genes in Holomycota. **a**, Binary heat-map and **b**, principal coordinate analysis (PCoA) species clustering based on the presence/absence of 1,158 orthologous genes belonging to 8 primary metabolism Gene Ontology categories across 43 eukaryotic genomes and transcriptomes. Species are color- and shaped-coded according to their taxonomic affiliation as indicated in the legend. COG presence is depicted in blue and absence is depicted in white.

At a more detailed level, the main differences in the metabolic complement of sanchytrids and of canonical fungi (+*Paraphelidium*) concerned the carbohydrate and lipid transport and metabolism categories, for which sanchytrids clustered with rozellids (Supplementary Fig. 4). We further pairwise-compared KEGG^71^ orthologs of sanchytrids against *R. allomycis* and the blastocladiomycete *A. macrogynus* (as representative of the sanchytrid closest canonical fungal relatives). The KEGG metabolic maps of *A. gromovi* and *S. tribonematis* contained 1,222 and 1,418 orthologous groups, respectively, whereas those of *R. allomycis* and *A. macrogynus* contained 845 and 4,860, respectively (Supplementary Fig. 5a-c). Blastocladiomycetes and sanchytrids shared more similarities, including the maintenance of amino acid and nucleotide metabolism and energy production with a complete electron transport chain, which were largely lost in *Rozella*^9,13^. Nonetheless, a reductive trend in energy production pathways could be observed in sanchytrid mitochondria, including the loss of ATP8, one F-type ATP synthase subunit that is also absent or highly modified in several metazoans, including chaetognaths, rotifers, most bivalve molluscs, and flatworms^72,73^. *S. tribonematis* also lacked the NADH dehydrogenase subunit NAD4L (Supplementary Fig. 1), although this loss is unlikely to impact its capacity to produce ATP since *R. allomycis*, which lacks not only ATP8 but also the complete NADH dehydrogenase complex, still seems to be able to synthesize ATP^9^.

Most carbohydrate-related metabolic pathways were retained in sanchytrids and canonical fungi except for the galactose and inositol phosphate pathways, absent in both sanchytrids and *Rozella*. Nonetheless, sanchytrids displayed a rich repertoire of carbohydrate-degrading enzymes (Supplementary Figs. 6-10), most of them being likely involved in the degradation of algal cell walls required for penetration into the host cells^13,74,75^. The most important difference with canonical fungi concerned lipid metabolism, with the steroids and fatty acid metabolism missing in sanchytrids and also in *Rozella*^9^ (Supplementary Fig. 5d-i). Collectively, our data suggest that, compared to Blastocladiomycota and other fungal relatives, sanchytrids have undergone a metabolic reduction that seems convergent with that observed in the phylogenetically distinct rozellid parasites.

### Convergent reductive flagellum evolution in Holomycota

The loss of the ancestral opisthokont single posterior flagellum in terrestrial fungi^76,77^ is thought to have been involved in their adaptation to land environments^78^. The number and timing of flagellum losses along the holomycotan branch remains to be solidly established. The flagellum is completely absent in nucleariids^12,52^ but is found in representatives of all other major holomycotan clades to the exclusion of the highly derived Microsporidia. They include rozellids^79^, aphelids^10^, and various canonical fungal groups, namely chytrids^70^, blastocladiomycetes^80^, *Olpidium*^21,26^, and sanchytrids^33,34,39^, although the latter are atypical. Sanchytrid amoeboid zoospores have never been observed swimming but gliding on solid surfaces via two types of pseudopods: thin filopodia growing in all directions and a broad hyaline pseudopodium at the anterior end (Fig. 1). The posterior flagellum, often described as a pseudocilium, drags behind the cell without being involved in active locomotion^33,34^. Its basal body (kinetosome) and axoneme ultrastructure differs from that of most flagellated eukaryotes. Instead of the canonical kinetosome with 9 microtubule triplets and axonemes with 9 peripheral doublets + 2 central microtubules^81^, sanchytrids exhibit reduced kinetosomes (9 singlets in *S. tribonematis*; 9 singlets or doublets in *A. gromovi*) and axonemes (without the central doublet and only 4 microtubular singlets)^33,34^. Despite this substantial structural simplification, sanchytrid kinetosomes are among the longest known in eukaryotes, up to 2.2 μm^33,34^. Such long but extremely simplified kinetosomes have not been reported in any other zoosporic fungi, including Blastocladiomycota^33,34^. Some of them, including *P. sedebokerense*^82^, display amoeboid zoospores during the vegetative cycle, flagellated cells being most likely gametes^83–85^.

To better understand flagellar reduction and loss across Holomycota, we analysed 61 flagellum-specific proteins on a well-distributed representation of 43 flagellated and non-flagellated species. Sanchytrids lacked several functional and maintenance flagellar components (Fig. 4a), namely axonemal dyneins, single- and double-headed inner arm dyneins, all intraflagellar transport proteins (IFT) of the group IFT-A and several of the group IFT-B. Sanchytrid kinetosomes have also lost several components of the centriolar structure and tubulins, including Centrin2, involved in basal body anchoring^86^, and Delta and Epsilon tubulins, essential for centriolar microtubule assembly and anchoring^87^. These losses (Fig. 4b) explain why sanchytrids lack motile flagella. Cluster analyses based on the presence/absence of flagellar components (Supplementary Fig. 11) showed sanchytrids at an intermediate position between flagellated and non-flagellated lineages. Therefore, sanchytrids are engaged in an unfinished process of flagellum loss, thereby providing an interesting model to study intermediate steps of this reductive process.

**Fig. 4.**
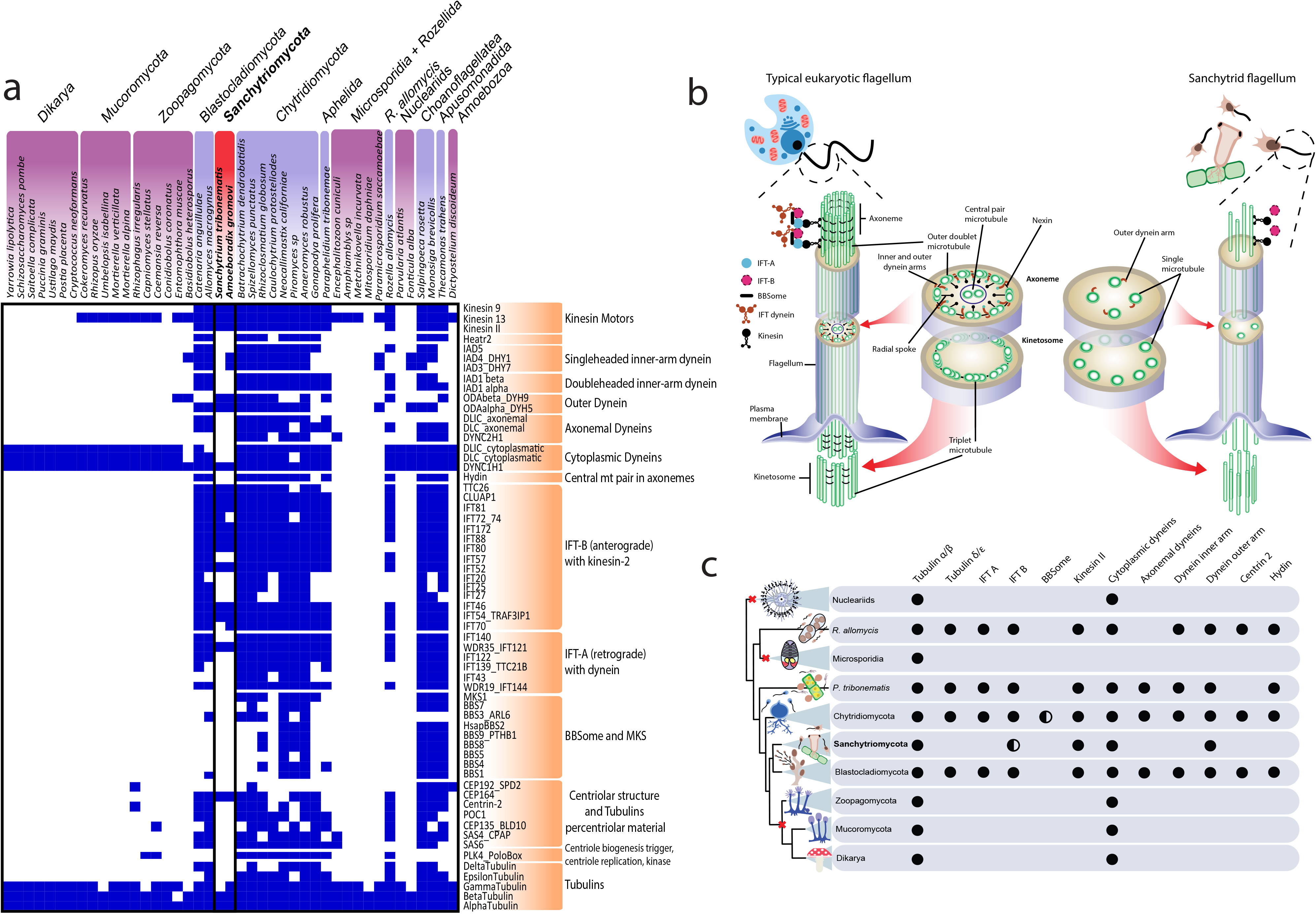
Comparison of flagellar protein distribution and structure in sanchytrids and other Holomycota. **a**, Presence/absence heatmap of 61 flagellum-specific proteins in 43 eukaryotic flagellated (purple; red: Sanchytriomycota) and non-flagellated (pink) lineages. The right column lists microtubular genes and flagellum-specific genes (orange). **b**, Illustration depicting the main structural elements present in a typical eukaryotic flagellum (left) and a reduced sanchytrid flagellum (right). Gene presence is depicted in blue and absence in white. **c**, Representation of the evolutionary relationships of holomycotan lineages and their patterns of presence/absence of key molecular components of the flagellar apparatus (full or half circles). Red crosses on branches indicate independent flagellum losses.

In addition to sanchytrid reduction, between four and six independent flagellar losses have been inferred in Holomycota^32^. Our new, more robust phylogenetic framework (Fig. 2) allowed to infer three large independent flagellum losses, plus the ongoing one in sanchytrids (Fig. 4c). These losses occurred at the base of high-rank taxa: nucleariids, Microsporidia, and the Zoopagomycota+Mucoromycota+Dikarya clade. A possible fourth loss event occurred in *Hyaloraphidium curvatum*, an atypical non-flagellated fungus originally classified as a colourless green alga^14^ and later reclassified within the Monoblepharidomycota^26,88^. Further analysis will be needed to confirm loss of flagellar components in this species. In addition, a putative fifth independent loss might have occurred in the Nephridiophagida, a clade of fungal parasites of insects and myriapods^89,90^ without clear affinity with established fungal clades^90^. Recently, a possible relationship to chytrids has been suggested^91^, though genomic or transcriptomic data to clarify their phylogenetic position are still missing.

### Fungal “vision” and flagellum exaptation

Why do sanchytrids retain a non-motile flagellum with a simplified but very long kinetosome? Since the primary flagellar function has been lost in favour of amoeboid movement, other selective forces must be acting to retain this atypical structure for a different function in zoospores. In bacteria, the exaptation of the flagellum for new roles in mechanosensitivity^92,93^ and wetness sensing^94^ has been documented. Microscopy observations of sanchytrid cultures showed that the flagellum is rather labile and can be totally retracted within the cell cytoplasm, the long kinetosome likely being involved in this retraction capability^33,34^. Interestingly, a conspicuous curved rosary chain of lipid globules has been observed near the kinetosome in *A. gromovi* zoospores, often also close to mitochondria^33,34^. In the blastocladiomycete *B. emersonii*, similar structures tightly associated with mitochondria are known as “side-body complexes”^95^. *B. emersonii* possesses a unique bacterial type-1-rhodopsin+guanylyl cyclase domain fusion (BeGC1, 626 amino acids) which, together with a cyclic nucleotide-gated channel (BeCNG1), controls zoospore phototaxis in response to cGMP levels after exposure to green light^96^. BeGC1 was localized by immunofluorescence on the external membrane of the axoneme-associated lipid droplets, which function as an eyespot at the base of the flagellum and control its beating^96–98^. The BeGC1 fusion and the channel BeCNG1 proteins have also been found in other blastocladiomycetes (*A. macrogynus* and *C. anguillulae*).

Both *A. gromovi* and *S. tribonematis* possessed the BeGC1 fusion (532 and 535 amino acids, respectively) and the gated channel BeCNG1 (Supplementary Fig. 12a-c). Therefore, this fusion constitutes a shared trait in Blastocladiomycota and Sanchytriomycota. Despite some ultrastructural differences and the need of functional studies to confirm their role, the presence of lipid threads in the vicinity of the kinetosome and mitochondria, together with the BeGC1 and BeCNG1 homologs, suggest the existence of a comparable light-sensing organelle in *Amoeboradix* and *Sanchytrium*. We hypothesize that, as in *B. emersonii*, the sanchytrid reduced flagellum could be involved in phototactic response, at least as a structural support for the lipid droplets. Interestingly, sanchytrids showed considerably shorter branches in rhodopsin and guanylyl cyclase domain phylogenetic trees (Supplementary Fig. 12a-b) than in multi-gene phylogenies (Fig. 2), indicating that these proteins (and their functions) are subjected to strong purifying selection as compared to other proteins encoded in their genomes.

Since rhodopsins capture light by using the chromophore retinal^99^, we looked for the carotenoid (β-carotene) biosynthesis enzymes^96,100^ necessary for retinal production. Surprisingly, the enzymes involved in the classical pathway (bifunctional lycopene cyclase/phytoene synthase, phytoene dehydrogenase and carotenoid oxygenase)^96,100^ were missing in both sanchytrid genomes, suggesting that they are not capable to synthesize their own retinal (Supplementary Data 5). We only detected two enzymes (isopentenyl diphosphate isomerase and farnesyl diphosphate synthase) that carry out early overlapping steps in biosynthesis of both sterol and carotenoids. By contrast, the β-carotene biosynthesis pathway is widely distributed in Fungi, including chytrids and blastocladiomycetes (*Allomyces* and *Blastocladiella*^96,101^). Therefore, sanchytrids, like most heterotrophic eukaryotes, seem unable to synthesize β-carotene and must obtain carotenoids through their diet^100^. Indeed, we detected all carotenoid and retinal biosynthesis genes in the transcriptome of the yellow-brown alga *Tribonema gayanum* (Supplementary Data 5), their host and likely retinal source during infection.

### Evolution of multicellularity in Holomycota

Fungal multicellularity results from connected hyphae^102^. Diverse genes involved in hyphal multicellularity were present in the ancestors of three lineages of unicellular fungi (Blastocladiomycota, Chytridiomycota and Zoopagomycota; BCZ nodes)^103^. To ascertain whether they were also present in other deep-branching Holomycota with unicellular members, we reconstructed the evolutionary history of 619 hyphal morphogenesis proteins^103^, grouped into 10 functional categories (see Methods; Supplementary Data 6; Supplementary Fig. 13). Our results showed that most hyphal morphogenesis genes were not only present in the last common fungal ancestor but also in other unicellular holomycotan relatives, indicating that they evolved well before the origin of fungal multicellularity (Fig. 5a). This pattern could be observed for all functional categories with the clear exception of the adhesion proteins, most of which only occur in Dikarya (Supplementary Fig. 13), reinforcing previous conclusions that adhesion proteins played a marginal role in the early evolution of hyphae^103^ but highlighting their crucial role in current (fruiting body-producing) hyphal-based multicellularity.

**Fig. 5.**
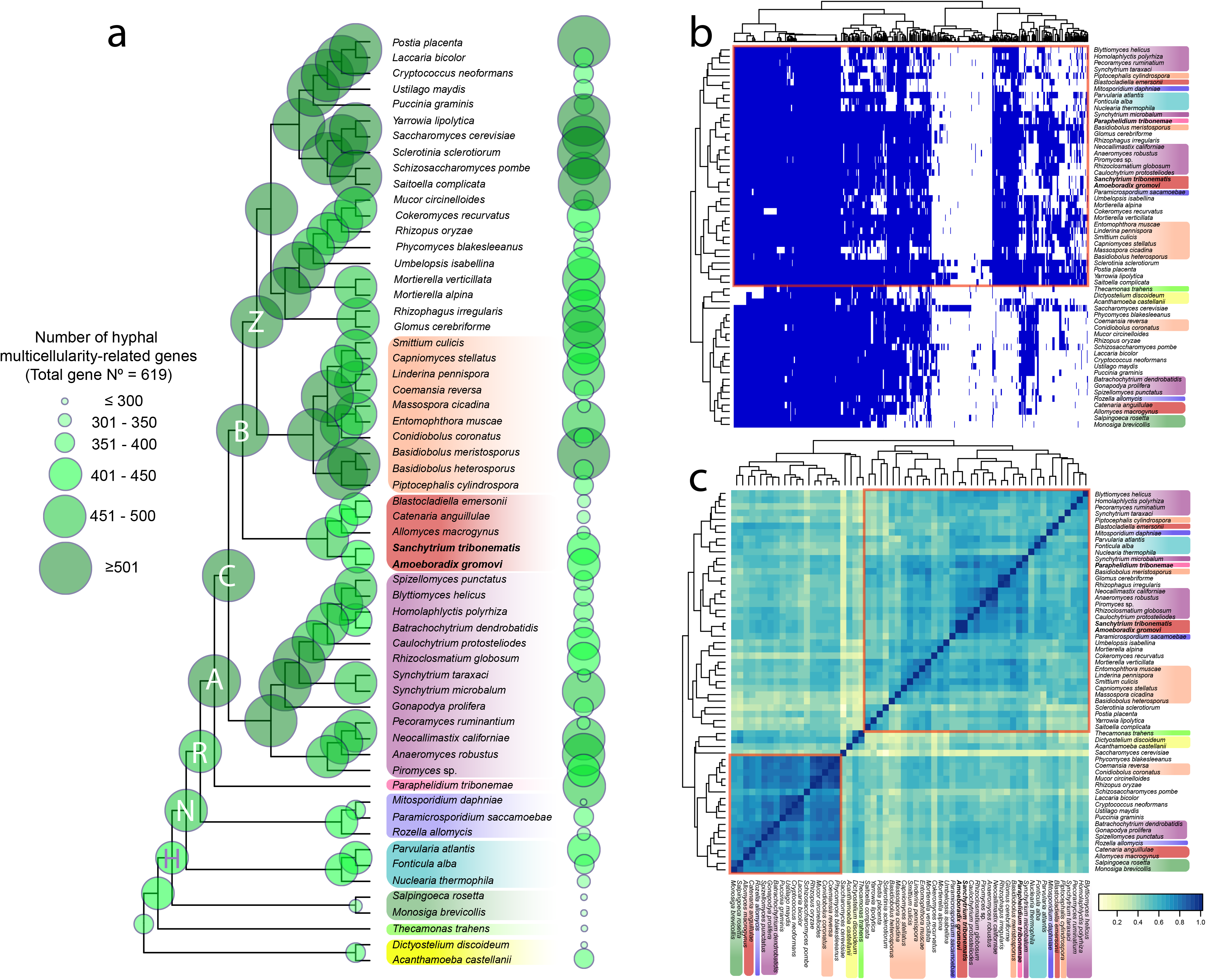
Hyphae-related genes across Holomycota. **a**, Cladogram of Holomycota depicting the phylogenetic relationships according to the GBE59 phylogenomic reconstruction. Bubble size on the nodes and tips represents the total number of reconstructed ancestral and extant hyphal multicellularity-related proteins. CBZ (Chytridiomycota, Blastocladiomycota and Zoopagomycota) and NRA (Nucleariids, Rozellida-Microsporidia and Aphelida) nodes are indicated with letters within the corresponding bubbles. Unicellular lineages are highlighted in colors according to their taxonomic affiliation (bottom to top: Amoebozoa [yellow], Apusomonadida [light green], Holozoa [dark green], nucleariids [light blue], Rozellida-Microsporidia [purple], Aphelida [pink], Chytridiomycota [violet], Sanchytriomycota-Blastocladiomycota [red] and Zoopagomycota [orange]). **b**, Presence/absence heatmap of 619 hyphal morphogenesis proteins in 59 unicellular (taxa are color-highlighted as in Fig. 5a) and multicellular (not color-highlighted) eukaryotic proteomes. Gene presence is depicted in blue and absence in white. **c**, Heatmap clustered by similarity showing the correlation between unicellular (taxa are color-highlighted as in Fig. 5a) and multicellular (not color-highlighted) taxa according to the presence/absence of hyphal morphogenesis gene proteins.

The common ancestor of sanchytrids and blastocladiomycetes possessed a high percentage of hyphae-related proteins (88.4%, node B in Fig. 5a), though this ancestral repertoire became secondarily reduced in sanchytrids (66.6%). Likewise, many yeasts, which are also secondarily reduced organisms, retained most of the genetic repertoire needed for hyphal development. Many of these proteins were also present in nucleariids, Rozellida-Microsporidia and Aphelida (nodes N, R, and A in Fig. 5a and Supplementary Data 7). Consequently, clustering analysis based on the presence/absence of hyphal morphogenesis proteins did not clearly segregate unicellular and multicellular lineages (Fig. 5b) and retrieved very weak intragroup correlation (Fig. 5c). Our results extend previous observations of hyphal morphogenesis genes from Fungi^103^ to much more ancient diversifications in the Holomycota. The holomycotan ancestor already possessed a rich repertoire of proteins, notably involved in ‘actin cytoskeleton’ and ‘microtubule-based transport’, that were later recruited for hyphal production (Supplementary Data 7). Most innovation concerned the proteins involved in the ‘cell wall biogenesis/remodeling’ and ‘transcriptional regulation’ functional categories, which expanded since the common ancestor of Aphelida and Fungi. This pattern is consistent with the enrichment of gene duplications in these two categories in all major fungal lineages^103^. Nevertheless, genome and transcriptome data remain very scarce for Aphelida and we expect that part of these duplications will be inferred to be older when more data for this sister lineage of Fungi become available.

## Conclusions

We generated the first genome sequence data for the two known species of sanchytrids, a group of atypical fungal parasites of algae. The phylogenetic analysis of two independent datasets of conserved proteins showed that they form a new fast-evolving fungal phylum, the Sanchytriomycota, sister to the Blastocladiomycota. Our phylogenetic analyses also provided strong support for Chytridiomycota being the sister group to all other fungi. Sanchytrids have a complex life cycle that includes a flagellated phase (zoospores) with non-motile flagella that are engaged in an ongoing reductive process. The inclusion of sanchytrids and a wide taxon sampling of fungi in our multi-gene phylogeny allowed the inference of three large independent flagellum loss events across Holomycota. Interestingly, the sanchytrid residual flagellum endowed with a long kinetosome might represent an exaptation of this structure as support for a lipid organelle probably involved in light sensing. Our taxon-rich dataset of deep-branching Holomycota also provided evidence for a very ancient origin of most genes related to hyphal morphogenesis, well before the evolution of the multicellular fungal lineages.

## Taxonomic appendix

### Sanchytriomycota phyl. nov.

Monocentric thallus, epibiotic; usually amoeboid zoospores with longest-known kinetosome in fungi (1-2 μm) and immobile pseudocilium; centrosome in sporangium with two centrioles composed by nine microtubular singlets.

Index Fungorum ID: IF558519

Class **Sanchytriomycetes** (Tedersoo et al. 2018)^104^ emend.

Diagnosis as for the phylum.

Order **Sanchytriales** (Tedersoo et al. 2018)^104^ emend.

Diagnosis as for the phylum.

Family **Sanchytriaceae** (Karpov & Aleoshin, 2017)^39^ emend.

Amoeboid zoospores with anterior lamellipodium producing subfilopodia, and lateral and posterior filopodia; with (rarely without) posterior pseudocilium; kinetosome composed of nine microtubule singlets or singlets/doublets, 1–2 μm in length. Zoospores attach to algal cell wall, encyst, and penetrate host wall with a short rhizoid. Interphase nuclei in sporangia have a centrosome of two centrioles composed of nine microtubular singlets. Predominantly parasites of freshwater algae.

Type: ***Sanchytrium*** (Karpov et Aleoshin 2017^39^) emend. Karpov, 2019^34^.

Parasite of algae. Epibiotic, spherical to ovate, sporangia with one (rarely more) discharge papillae. Amoeboid zoospores with anterior lamellipodium; with (rarely without) pseudocilium; contains kinetosome composed of nine single microtubules 1–1.2 μm in length. Interphase nuclei in sporangia have a centrosome with two orthogonal centrioles composed of nine microtubular singlets and with an internal fibrillar ring.

Type: *Sanchytrium tribonematis* (Karpov et Aleoshin, 2017^39^) emend. Karpov, 2019^34^.

***Sanchytrium tribonematis*** (Karpov et Aleoshin, 2017^39^) emend. Karpov, 2019^34^.

Round to ovate smooth sporangium, ~10 μm diameter, without or with one discharge papilla; sessile on algal surface. Slightly branched rhizoid, almost invisible inside host. Amoeboid zoospores 5.4 - 3.3 μm (maximum) with anterior lamellipodium producing subfilopodia, lateral and posterior filopodia; normally with posterior pseudocilium up to 5 μm in length supported by up to four microtubules.

***Amoeboradix*** Karpov, López-García, Mamkaeva & Moreira, 2018^33^.

Zoosporic fungus with monocentric, epibiotic sporangia and amoeboid zoospores having posterior pseudocilium that emerges from long kinetosome (ca. 2 μm) composed of microtubular singlets or doublets.

Type species: *A. gromovi* Karpov, López-García, Mamkaeva & Moreira, 2018^33^.

***A. gromovi*** Karpov, López-García, Mamkaeva & Moreira, 2018^33^.

Amoeboid zoospores of 2-3×3-4 μm with posterior pseudocilium up to 8 μm in length, which can be totally reduced; kinetosome length, 1.8-2.2 μm. Sporangia of variable shape, from pear-shaped with rounded broad distal end and pointed proximal end to curved asymmetrical sac-like 8-10 μm width 16-18 μm long with prominent rhizoid and 2-3 papillae for zoospore discharge; spherical cysts about 3-4 μm in diameter; resting spore ovate and thick-walled 12×6 μm. Parasite of *Tribonema gayanum, T. vulgare* and *Ulothrix tenerrima.*

## Methods

### Biological material

*Sanchytrium tribonematis* strain X-128 and *Amoeboradix gromovi* strain X-113, isolated from freshwater sampling locations in Russia^33,34^, were maintained in culture with the freshwater yellow-green alga *Tribonema gayanum* Pasch. strain 20 CALU as host ^39^. The algal host was grown in mineral freshwater medium at room temperature under white light. After inoculation with *Sanchytrium* or *Amoeboradix*, cultures were incubated for 2 weeks to reach the maximum infection level. We then collected both individual zoospores and sporangia full of moving zoospores by micromanipulation with an Eppendorf PatchMan NP2 micromanipulator using 19 μm VacuTip microcapillaries (Eppendorf) on an inverted Leica Dlll3000 B microscope. Sporangia were separated from the algal host cells using a microblade mounted on the micromanipulator. Zoospores and sporangia were washed 2 times in clean sterile water drops before storing them into individual tubes for further analyses.

### Whole genome amplification and sequencing

DNA extraction from single zoospores and sporangia was done with the PicoPure kit (Thermo Fisher Scientific) according to the manufacturer’s protocol. Whole genome amplification (WGA) was carried out by multiple displacement amplification with the single-cell REPLI-g kit (QIAGEN). DNA amplification was quantified using a Qubit fluorometer (Life Technologies). We retained WGA products that yielded high DNA concentration. As expected, WGA from sporangia (many zoospores per sporangium) yielded more DNA than individual zoospores and were selected for sequencing (K1-9_WGA for *A. gromovi*; SC-2_WGA for *S. tribonematis*). TruSeq paired-end single-cell libraries were prepared from these samples and sequenced on a HiSeq 2500 Illumina instrument (2 x 100 bp) with v4 chemistry. We obtained 121,233,342 reads (26,245 Mbp) for *A. gromovi* and 106,922,235 reads (21,384 Mbp) for *S. tribonematis*.

### Genome sequence assembly, decontamination and annotation

Paired-end read quality was assessed with FastQC v0.11.9^105^ before and after quality trimming. Illumina adapters were removed with Trimmomatic v0.32 in Paired-End mode^106^, with the following parameters: ILLUMINACLIP:adapters.fasta:2:30:10LEADING:28 TRAILING:28 SLIDINGWINDOW:4:30. Trimmed paired-end reads were assembled using SPAdes v3.9.1 in single-cell mode^107^. This produced assemblies of 48.7 and 37.1 Mb with 8,420 and 8,015 contigs for *A. gromovi* and *S. tribonematis*, respectively. Elimination of contaminant contigs was carried out by a three-step process. First, genome sequences were subjected to two rounds of assembly, before and after bacterial sequence removal with BlobTools v0.9.19^108^ (Supplementary Fig. 14). Second, open-reading frames were predicted and translated from the assembled contigs using Transdecoder v2 (http:transdecoder.github.io) with default parameters and Cd-hit v4.6^109^ with 100% identity to produce protein sequences for *A. gromovi* and *S. tribonematis*. Finally, to remove possible eukaryotic host (*Tribonema*) contamination, the predicted protein sequences were searched by BLASTp^110^ against two predicted yellow-green algae proteomes inferred from the *Tribonema gayanum* transcriptome ^13^ and the *Heterococcus* sp. DN1 genome (PRJNA210954; also a member of the Tribonematales). We excluded sanchytrid hits that were 100% or >95% identical to them, respectively. Statistics of the final assembled genomes were assessed with QUAST v4.5^111^ and Qualimap v2.2.1^112^ for coverage estimation. In total, we obtained 7,220 and 9,368 protein sequences for *A. gromovi* and *S. tribonematis*, respectively (Table 1). To assess genome completeness, we used BUSCO v2.0.1^37^ on the decontaminated predicted proteomes with the fungi_odb9 dataset of 290 near-universal single-copy orthologs. The inferred proteins were functionally annotated with eggNOG mapper v2^38^ using DIAMOND as the mapping mode and the eukaryotic taxonomic scope. This resulted in 3,757 (*A. gromovi*) and 4,670 (*S. tribonematis*) functionally annotated peptides for the predicted proteomes (Supplementary Data 1). To determine the presence of sanchytrid-specific proteins we used OrthoFinder^43^ to generate orthologous groups using the proteomes of the 59 species used in the GBE59 dataset (Supplementary Data 2). After extracting the orthogroups that were present only in both sanchytrid species, we functionally annotated them with eggNOG mapper v2^38^ (Supplementary Data 2). The mitochondrial genomes of the two sanchytrid species were identified in single SPAdes-generated contigs using BLAST^113^ and annotated with MITOS v1^114^. We confirmed the quality of the obtained mitochondrial genome assemblies by reassembling them with NOVOPlasty^115^. Additional BLAST searches were made to verify missing proteins.

### Phylogenomic analyses and single-gene phylogenies

A revised version of a 264 protein dataset (termed GBE)^49^ was used to reconstruct phylogenomic trees^13,52^. This dataset was updated with sequences from the two sanchytrid genomes and all publicly available Blastocladiomycota sequences. Sequences were obtained mainly from GenBank (http://www.ncbi.nlm.nih.gov/genbank, last accessed November, 2019) and, secondarily, the Joint Genome Institute (http://www.jgi.doe.gov/; last accessed May 2017). For details on the origin of sequence data see Supplementary Data 8. Our updated taxon sampling comprises a total of 71 Opisthokonta (2 Holozoa and 69 Holomycota), 2 Amoebozoa and 1 Apusomonadida. Two datasets with two different taxon samplings were prepared, one with all 74 species (GBE74) and one without the long-branching core Microsporidia and metchnikovellids for a total of 59 species (GBE59). In addition, these datasets were also constructed without the sanchytrids to test if they induced LBA artefacts, for a total of 72 (GBE 72) and 57 (GBE57) species. Additionally, for the taxon sampling of 59 species we used a second phylogenomic dataset of 53 highly conserved proteins^59^; the BMC dataset (BMC59).

The proteins for these datasets were searched in the new species with tBLASTn^110^, incorporated into the individual protein datasets, aligned with MAFFT v7 ^116^ and trimmed with TrimAl v1.2 with the automated1 option^117^. Alignments were visualized, manually edited and concatenated with Geneious v6.0.6^118^ and single-gene trees obtained with FastTree v2.1.7^119^ with default parameters. Single-gene trees were manually checked to identify and remove paralogous and/or contaminating sequences. The concatenation of the clean trimmed 264 proteins of the GBE dataset resulted in alignments containing 93,743 (GBE59) and 86,313 (GBE74) amino acid positions. The alternative alignment without the sanchytrids contained 93,421 (GBE57) and 84,949 (GBE72) amino acid positions. The concatenation of the 53 proteins of the BMC59 dataset resulted in an alignment of 14,965 amino acid positions. The Bayesian inference (BI) phylogenetic trees of the GBE59 and BMC59 datasets were reconstructed using PhyloBayes-MPI v1.5^120^ under the CAT-Poisson and CAT-GTR models, respectively, with two MCMC chains and run for more than 15,000 generations, saving one every 10 trees. Analyses were stopped once convergence thresholds were reached (i.e. maximum discrepancy <0.1 and minimum effective size >100, calculated using bpcomp) and consensus trees constructed after a burn-in of 25%. Maximum likelihood (ML) phylogenetic trees were inferred with IQ-TREE v1.6 under the PMSF approximation of the LG+C20+R8 (GBE59 and GBE57), LG+C20+F+R9 (GBE72), LG+C20+F+R10 (GBE74) or LG+C60+R7 (BMC59) models with guide trees inferred with the LG+R8, LG+F+R9, LG+F+R10 or LG+R7 models, respectively, selected with the IQ-TREE TESTNEW algorithm as per the Bayesian information criterion (BIC). Statistical support was generated with 1,000 ultrafast bootstraps^121^ and 1,000 replicates of the SH-like approximate likelihood ratio test^122^. All trees were visualized with FigTree^123^. Additionally, we reconstructed an ML tree for BMC59 under the same model with 100 conventional bootstrap replicates.

To test alternative tree topologies we used Mesquite^124^ to constrain the following topologies: 1) chytrids as sister lineage of all other fungi (i.e., monophyly of Blastocladiomycota+Sanchytriomycota+Zygomycota+Dikarya), and 2) Blastocladiomycota+Sanchytriomycota as sister lineage of all other fungi (i.e., monophyly of Chytridiomycota+Zygomycota+Dikarya). The constrained topologies without branch lengths were reanalysed with the -g option of IQ-TREE and the best-fitting model. AU tests were carried out on the resulting trees for each taxon sampling with the -z and -au options of IQ-TREE. Additionally, to minimize possible systematic bias due to the inclusion of fast-evolving sites in our GBE59 dataset, we progressively removed the fastest evolving sites, 5% of sites at a time. For that, among-site substitution rates were inferred using IQ-TREE under the -wsr option and the best-fitting model for a total of 19 new data subsets (Supplementary Data 3). We then reconstructed phylogenetic trees for all these subsets using IQ-TREE with the same best-fitting model as for the whole dataset. To assess the support of the alternative topologies in the bootstrapped trees, we used CONSENSE from the PHYLIP v3.695 package^125^ and interrogated the UFBOOT file using a Python script (M. Kolisko, pers. comm). Single-protein phylogenies were reconstructed by ML using IQ-TREE with automatic best fit model selection.

### Comparative analysis of primary metabolism

We carried out statistical multivariate analyses to get insights into the metabolic capabilities of sanchytrids in comparison with other Holomycota. We searched in both sanchytrids 1,206 eggNOG orthologous groups^38^ corresponding to 8 primary metabolism categories (Gene Ontology, GO). The correspondence between GO terms and primary metabolism COGs used are the following: [C] Energy production and conversion (227 orthologs); [G] Carbohydrate transport and metabolism (205 orthologs); [E] Amino acid transport and metabolism (200 orthologs); [F] Nucleotide transport and metabolism (87 orthologs); [H] Coenzyme transport and metabolism (94 orthologs); [I] Lipid transport and metabolism (201 orthologs); [P] Inorganic ion transport and metabolism (153 orthologs); and [Q] Secondary metabolites biosynthesis, transport and catabolism (70 orthologs). From these categories, we identified 1,158 orthologs (non-redundant among categories) in the sanchytrid genomes which were shared among a set of 43 species, including 38 Holomycota, 2 Holozoa, 2 Amoebozoa, and 1 apusomonad (for the complete list, see Supplementary Data 4). We annotated the protein sets of these 43 species using eggNOG-mapper v2^38^ with DIAMOND as mapping mode and the eukaryotic taxonomic scope. All ortholog counts were transformed into a presence/absence matrix (encoded as 0/1) and analysed with the R script^126^ detailed in Torruella et al. (2018) in which similarity values between binary COG profiles of all species were calculated to create a complementary species-distance matrix. We then analysed this distance matrix using a Principal Coordinate Analysis (PCoA) and plotted binary COG profiles in a presence/absence heatmap. Clustering of the species and orthologs was done by Ward hierarchical clustering (on Euclidean distances for orthologs) of the interspecific Pearson correlation coefficients. The raw species clustering was also represented in a separate pairwise correlation heatmap colour-coded to display positive Pearson correlation values (0–1). Finally, COG categories of each primary metabolism were also analysed separately (categories C, E, F, G, H, I, P, and Q) using the same workflow (for more details see Torruella et al. 2018 and https://github.com/xgrau/paraphelidium2018). We compared in more detail the inferred metabolism of a subset of species (the two sanchytrids, *Rozella allomycis*, and *Allomyces macrogynus*). The annotation of these proteomes was done using BlastKOALA v2.2^127^, with eukaryotes as taxonomy group and the genus_eukaryotes KEGG GENES database, and the annotations were uploaded in the KEGG Mapper Reconstruct Pathway platform^71^ by pairs. First, we compared the two sanchytrid proteomes to confirm their similarity and, second, we compared them with the proteomes of *R. allomycis* and *A. macrogynus* to study their metabolic reduction. To explore if the long branch observed in sanchytrids could be explained by the absence of DNA repair and genome maintenance genes, we used a dataset of 47 proteins involved on these functions from a previous study on fast-evolving budding yeasts^60^. These proteins where blasted^113^ against the proteomes of the two sanchytrid species, the blastoclad *A. macrogynus* and four species of fast- and slow-evolving yeasts from that study.

### Homology searches and phylogenetic analysis of specific proteins

To assess the evolution of the flagellum in holomycotan lineages we used a dataset of over 60 flagellum-specific proteins ^77,128–130^ to examine a total of 43 flagellated and non-flagellated species within and outside the Holomycota. The flagellar toolkit proteins were identified using *Homo sapiens* sequences as Blast queries. Candidate proteins were then blasted against the non-redundant GenBank database to confirm their identification and submitted to phylogenetic analysis by multiple sequence alignment with MAFFT^116^, trimming with TrimAl^117^ with the automated1 option, and tree reconstruction with FastTree^119^. After inspection of trees, we removed paralogs and other non-orthologous protein sequences. We excluded the proteins with no identifiable presence in any of the 43 species used in the analysis and encoded the presence/absence of the remaining ones in a 1/0 matrix. The native R heatmap function^126^ was used to plot the flagellar proteome comparison between all species according to their presence/absence similarity profiles. To study the presence or absence of the fusion of the BeGC1 and BeCNG1 light-sensing proteins, we blasted them against the proteomes of *S. tribonematis* and *A. gromovi* using the *Blastocladiella emersonii* sequences (BeGC1: AIC07007.1; BeCNG1: AIC07008.1) as queries. We used the protein dataset of Avelar et al. (2014)^96^ in which the authors blasted both the guanylyl-cyclase GC1 and the rhodopsin domains of the BeGC1 fusion protein of *B. emersonii* and the BeCNG1 proteins against a database of more than 900 genomes from eukaryotic and prokaryotic species all across the tree of life (see Table S1 of their manuscript). We used MAFFT to include the new sequences in a multiple sequence alignment for the BeCNG1 protein channel and separately for both domains of the BeGC1 fusion protein^96^. After trimming with TrimAl we reconstructed phylogenetic trees for the three datasets using IQ-TREE with the best-fitting models: LG+F+I+G4 for the rhodopsin domain and BeCNG1, and LG+G4 model for GC1 guanylyl cyclase domain. The resulting trees were visualized with FigTree^123^. To study the possible presence of a carotenoid synthesis pathway for retinal production in sanchytrids, we combined previous datasets for carotenoid biosynthesis and cleavage enzymes from one giant virus (ChoanoV1), two choanoflagellates and two haptophytes^100^, together with the carotenoid biosynthesis enzymes found in *B. emersonii*^96^. This dataset also included three enzymes for early sterol and carotenoid biosynthesis (isoprenoid biosynthesis steps). The protein sequences from this dataset where blasted (BLASTp) against the proteomes of the two sanchytrids and their host *Tribonema gayanum*. Results were confirmed by blasting the identified hits to the NCBI non-redundant protein sequence database. To study the presence of proteins involved in cell-wall penetration in sanchytrids (including cellulases, hemicellulases, chitinases, and pectinases), we used the complete mycoCLAP v1 database of carbohydrate-degrading enzymes (https://mycoclap.fungalgenomics.ca/mycoCLAP/). We blasted this database against our two sanchytrid proteomes and other two representatives of Blastocladiomycota and four Chytridiomycota. We then screened for canonical cellulose, hemicellulose, pectin and chitin degrading enzymes identified in Fungi from previous studies^13,131,132^. The identified hits were blasted back to both the NCBI non-redundant protein sequence database and the CAZy database (cazy.org) to create a dataset for phylogenetic reconstruction for each enzyme. We added the identified sequences to specific protein datasets from previous studies in the case of cellulases^13^ and of the chitin degradation proteins GH20 β -N-acetylhexosaminidase (NAGase)^75^ and GH18 chitinase^74^. Proteins were aligned with MAFFT^116^ and trimmed from gaps and ambiguously aligned sites using TrimAl^117^ with the automated1 option. ML trees were inferred using IQ-TREE^133^ with the best-fitting model selected with the IQ-TREE TESTNEW algorithm as per BIC. The best-scoring tree was searched for up to 100 iterations, starting from 100 initial parsimonious trees; statistical supports were generated from 1000 ultra-fast bootstrap replicates and 1000 replicates of the SH-like approximate likelihood ratio test. Trees were visualized with FigTree^123^.

### Analyses of hyphae multicellularity-related genes

We used a dataset of 619 hyphal multicellularity-related proteins belonging to 362 gene families^103^. These gene families were grouped into 10 functional categories: actin cytoskeleton (55 proteins), adhesion (30), polarity maintenance (107), cell wall biogenesis/remodeling (92), septation (56), signaling (82), transcriptional regulation (51), vesicle transport (103), microtubule-based transport (32) and cell cycle regulation (11). The 619 proteins were searched by BLAST in the GBE59 dataset proteome and later incorporated into individual gene family protein alignments with MAFFT v7^116^. After trimming with TrimAl^117^ with the automated1 option, alignments were visualized with Geneious v6.0.6^118^ and single gene trees obtained with FastTree^119^ with default parameters. Single protein trees were manually checked to identify paralogous sequences and confirm the presence of genes within large gene families. Presence/absence of genes was binary coded (0/1) for each species and the resulting matrix processed with a specific R script^13^. Finally, we reconstructed the multicellularity-related gene gain/loss dynamics by applying the Dollo parsimony method implemented in Count v10.04^134^. These analyses were done on both the complete dataset of all hyphae multicellularity-related genes and on 10 separate datasets corresponding to the functional categories mentioned above.

## Data availability

The raw sequence data and assembled genomes generated in this study have been deposited at the National Center for Biotechnology Information (NCBI) sequence databases under Bioproject accession codes PRJNA668693 [https://www.ncbi.nlm.nih.gov/bioproject/PRJNA668693] and PRJNA668694 [https://www.ncbi.nlm.nih.gov/bioproject/PRJNA668694]. Additional data generated in this study (including alignments, phylogenetic trees and assembled genomes) are available in the Figshare repository project 91439 [https://figshare.com/projects/Sanchytriomycota_Galindo_etal/91439]. DNA and protein sequences of the species used in this study were downloaded from the GenBank public databases nr [https://www.ncbi.nlm.nih.gov/nucleotide/ and https://www.ncbi.nlm.nih.gov/protein/], genome [https://www.ncbi.nlm.nih.gov/genome/], SRA [https://www.ncbi.nlm.nih.gov/sra/], the JGI genome database [https://genome.jgi.doe.gov/portal/], the CAZy database [cazy.org], and the mycoCLAP database [https://mycoclap.fungalgenomics.ca/mycoCLAP/], for more details see Supplementary Data 8.

## Acknowledgments

We thank the UNICELL single-cell genomics platform (https://www.deemteam.fr/en/unicell). This work was funded by the European Research Council Advanced Grants ProtistWorld (No. 322669, P.L.-G.) and Plast-Evol (No. 787904, D.M.) and the Horizon 2020 research and innovation program under the Marie Skłodowska-Curie ITN project SINGEK (http://www.singek.eu/; grant agreement No. H2020-MSCA-ITN-2015-675752, L.J.G. & P.L.-G.). S.A.K. contribution was supported by the RSF grant No. 16-14-10302. We thank Christina Cuomo and the Broad Institute for allowing the use of unpublished fungal genome sequences in this study.

## Author contributions

P.L.-G. and D.M. conceived and supervised the study. S.A.K. and D.M. prepared the biological material; L.J.G., G.T. and D.M. analysed the sequence data; L.J.G., P.L.-G. and D.M. supervised the study and wrote the manuscript, which was edited and approved by all authors.

**Correspondence and requests for materials** should be addressed to L.J.G or D.M.

