## Supplementary information for "Phylogenomics of a new fungal phylum reveals multiple waves of reductive evolution across Holomycota"

a

b

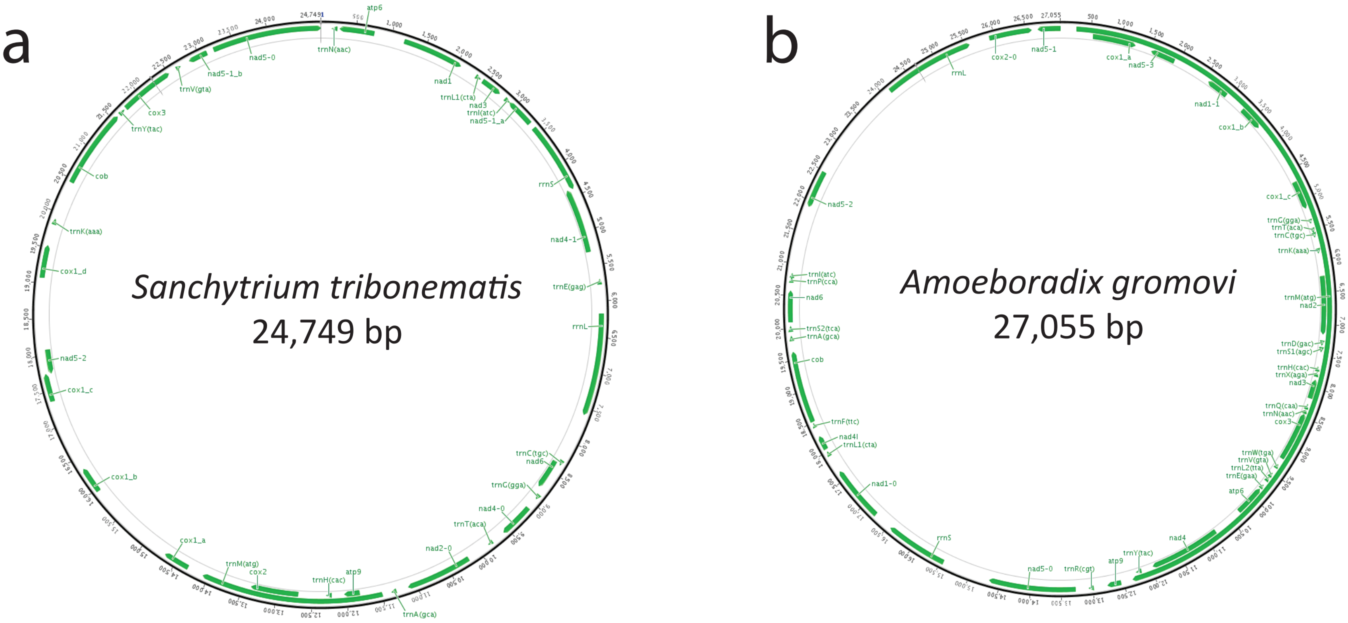


c

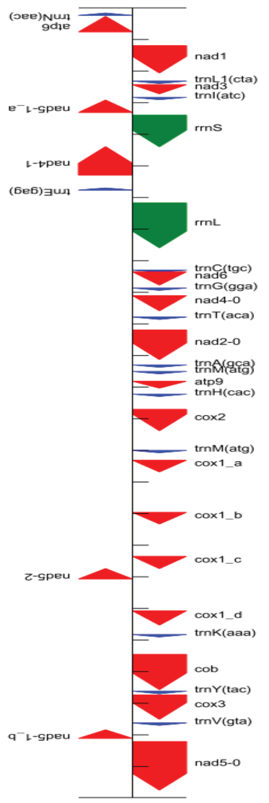


| Name | Start | Stop | Strand | Length | Structure |
|---|---|---|---|---|---|
| trnN(aac) | 167 | 239 | - | 73 | svg ps |
| atp6 | 271 | 762 | - | 492 | |
| nad1 | 1196 | 2062 | + | 867 | |
| trnL1(cta) | 2317 | 2391 | + | 75 | svg ps |
| nad3 | 2431 | 2721 | + | 291 | |
| trnI(atc) | 2832 | 2905 | + | 74 | svg ps |
| nad5-1_a | 2918 | 3310 | - | 393 | |
| rrnS | 3395 | 4397 | + | 1003 | svg ps |
| nad4-1 | 4392 | 5294 | - | 903 | |
| trnE(gag) | 5700 | 5765 | - | 66 | svg ps |
| rrnL | 6173 | 7593 | + | 1421 | svg ps |
| trnC(tgc) | 8296 | 8367 | + | 72 | svg ps |
| nad6 | 8352 | 8771 | + | 420 | |
| trnG(gga) | 8865 | 8936 | + | 72 | svg ps |
| nad4-0 | 9104 | 9586 | + | 483 | |
| trnT(aca) | 9783 | 9856 | + | 74 | svg ps |
| nad2-0 | 10191 | 11129 | + | 939 | |
| trnA(gca) | 11299 | 11371 | + | 73 | svg ps |
| trnM(atg) | 11505 | 11578 | + | 74 | svg ps |
| atp9 | 11814 | 12032 | + | 219 | |
| trnH(cac) | 12223 | 12293 | + | 71 | svg ps |
| cox2 | 12703 | 13380 | + | 678 | |
| trnM(atg) | 14003 | 14075 | + | 73 | svg ps |
| cox1_a | 14312 | 14683 | + | 372 | |
| cox1_b | 15964 | 16329 | + | 366 | |
| cox1_c | 17355 | 17741 | + | 387 | |
| nad5-2 | 17741 | 18070 | - | 330 | |
| cox1_d | 19076 | 19528 | + | 453 | |
| trnK(aaa) | 19830 | 19903 | + | 74 | svg ps |
| cob | 20456 | 21571 | + | 1116 | |
| trnY(tac) | 21618 | 21702 | + | 85 | svg ps |
| cox3 | 21737 | 22510 | + | 774 | |
| trnV(gta) | 22626 | 22698 | + | 73 | svg ps |
| nad5-1_b | 22843 | 23112 | - | 270 | |
| nad5-0 | 23214 | 24749 | + | 1536 | |

tRNA gene rRNA gene protein coding gene

d

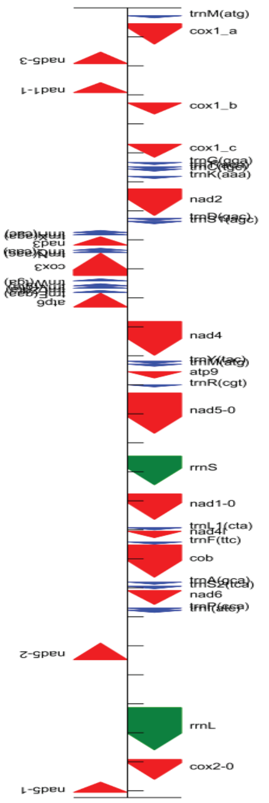


| Name | Start | Stop | Strand | Length | Structure |
|---|---|---|---|---|---|
| trnM(atg) | 272 | 344 | + | 73 | svg ps |
| cox1_a | 546 | 1253 | + | 708 | |
| nad5-3 | 1525 | 1923 | - | 399 | |
| nad1-1 | 2578 | 2925 | - | 348 | |
| cox1_b | 3280 | 3663 | + | 384 | |
| cox1_c | 4702 | 5154 | + | 453 | |
| trnG(gga) | 5331 | 5401 | + | 71 | svg ps |
| trnT(aca) | 5481 | 5555 | + | 75 | svg ps |
| trnC(tgc) | 5558 | 5630 | + | 73 | svg ps |
| trnK(aaa) | 5811 | 5884 | + | 74 | svg ps |
| nad2 | 6246 | 7163 | + | 918 | |
| trnD(gac) | 7262 | 7333 | + | 72 | svg ps |
| trnS1(agc) | 7364 | 7445 | + | 82 | svg ps |
| trnH(cac) | 7678 | 7749 | - | 72 | svg ps |
| trnX(aga) | 7755 | 7829 | - | 75 | svg ps |
| nad3 | 7895 | 8188 | - | 294 | |
| trnQ(caa) | 8294 | 8364 | - | 71 | svg ps |
| trnN(aac) | 8370 | 8441 | - | 72 | svg ps |
| cox3 | 8458 | 9231 | - | 774 | |
| trnW(tga) | 9349 | 9419 | - | 71 | svg ps |
| trnV(gta) | 9517 | 9588 | - | 72 | svg ps |
| trnL2(tta) | 9598 | 9672 | - | 75 | svg ps |
| trnE(gaa) | 9747 | 9818 | - | 72 | svg ps |
| atp6 | 9838 | 10320 | - | 483 | |
| nad4 | 10817 | 11983 | + | 1167 | |
| trnY(tac) | 12185 | 12267 | + | 83 | svg ps |
| trnM(atg) | 12288 | 12359 | + | 72 | svg ps |
| atp9 | 12545 | 12766 | + | 222 | |
| trnR(cgt) | 12999 | 13070 | + | 72 | svg ps |
| nad5-0 | 13285 | 14673 | + | 1389 | |
| rrnS | 15459 | 16450 | + | 992 | svg ps |
| nad1-0 | 16762 | 17643 | + | 882 | |
| trnL1(cta) | 17929 | 18002 | + | 74 | svg ps |
| nad4l | 18051 | 18272 | + | 222 | |
| trnF(ttc) | 18420 | 18492 | + | 73 | svg ps |
| cob | 18522 | 19637 | + | 1116 | |
| trnA(gca) | 19804 | 19877 | + | 74 | svg ps |
| trnS2(tca) | 19948 | 20031 | + | 84 | svg ps |
| nad6 | 20096 | 20575 | + | 480 | |
| trnP(cca) | 20702 | 20774 | + | 73 | svg ps |
| trnI(atc) | 20777 | 20849 | + | 73 | svg ps |
| nad5-2 | 21907 | 22482 | - | 576 | |
| rrnL | 24132 | 25591 | + | 1460 | svg ps |
| cox2-0 | 25923 | 26609 | + | 687 | |
| nad5-1 | 26702 | 27055 | - | 354 | |

tRNA gene rRNA gene protein coding gene

**Supplementary Fig. 1.** MITOS Graphical representation of the mitochondrial genome and gene content of *Sanchytrium tribonematis* (**a** and **c**), and *Amoeboradix gromovi* (**b** and **d**).

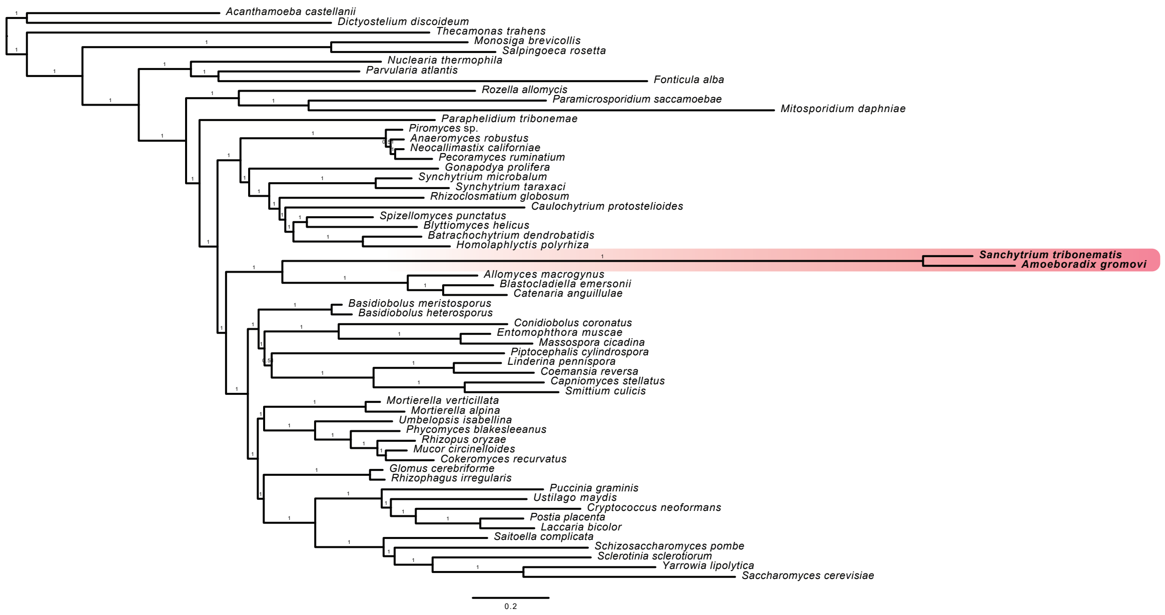


**Supplementary Fig. 2. a, Bayesian phylogenomic tree based on the GBE protein dataset.** The tree was reconstructed using 264 conserved proteins, 59 species, and 93,743 conserved amino acid positions, it was inferred using PhyloBayes under the CAT-Poisson model with posterior probability as statistical support.

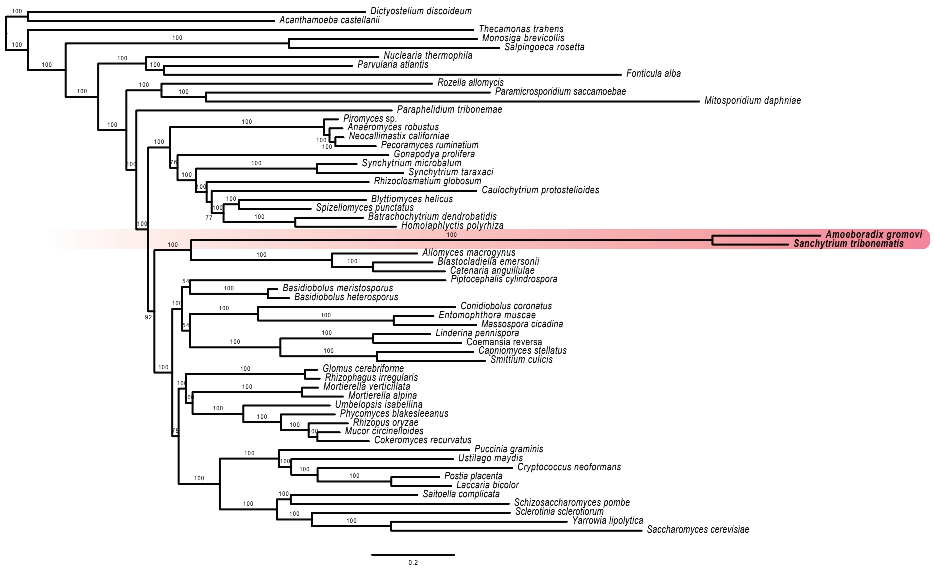


**Supplementary Fig. 2. b, Maximum likelihood tree based on the GBE protein dataset.** The tree was reconstructed using 264 conserved proteins, 59 species, and 93,743 conserved amino acid positions, it was inferred with IQ-TREE under the PMSF approximation of the LG+R8+C20 model and ultrafast bootstrap as statistical support.

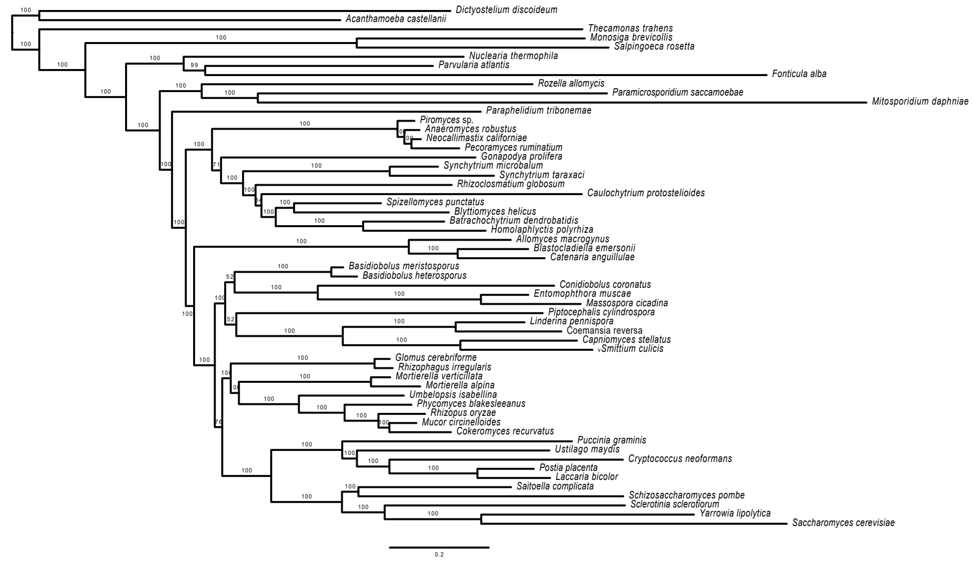


**Supplementary Fig. 2. c, Maximum likelihood tree based on the GBE protein dataset.** The tree was reconstructed using 264 conserved proteins, 57 species, and 93,421 conserved amino acid positions, it was inferred with IQ-TREE under the PMSF approximation of the LG+R8+C20 model and ultrafast bootstrap as statistical support.

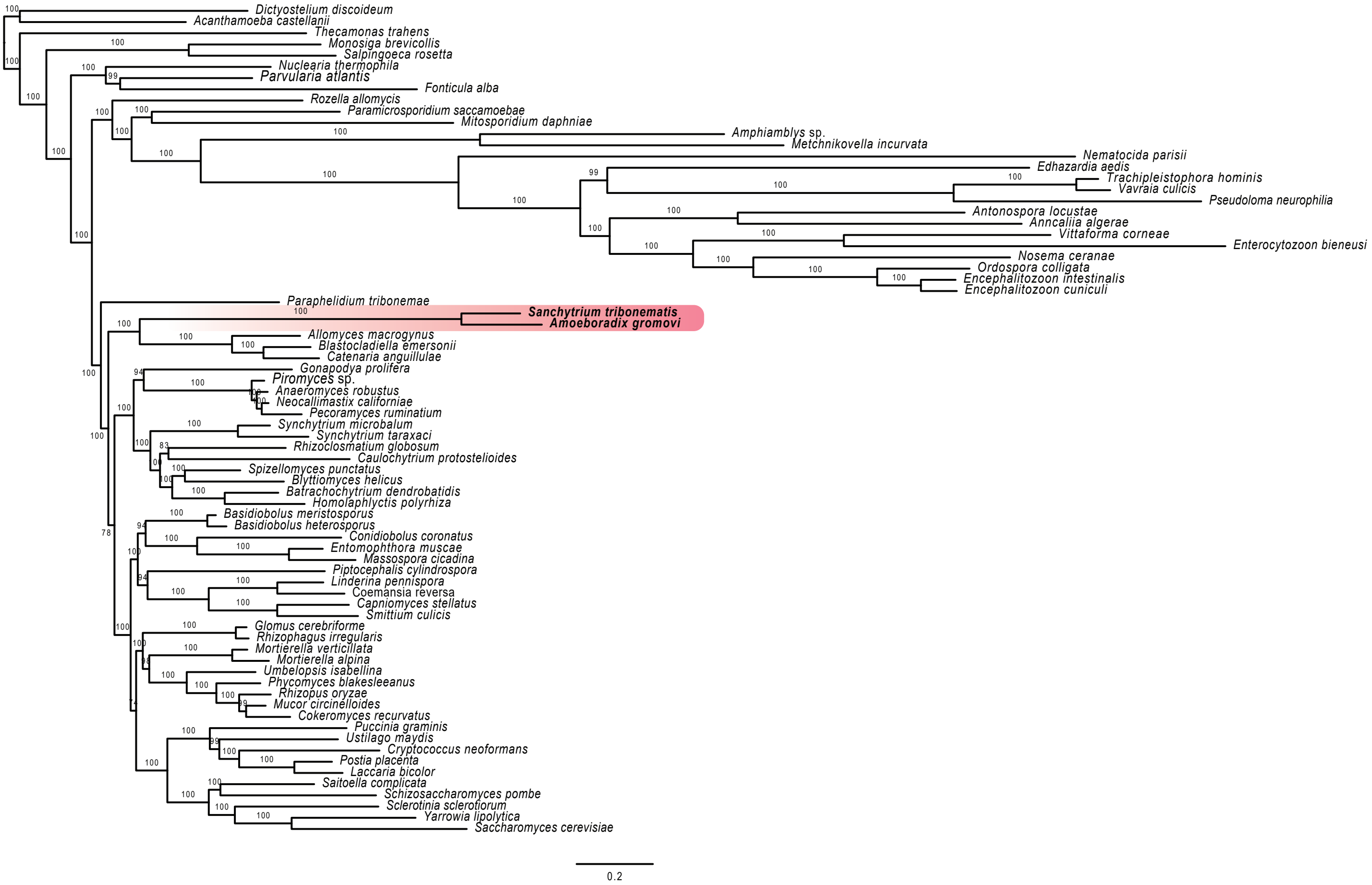


**Supplementary Fig. 2. d, Maximum likelihood tree based on the GBE protein dataset.** The tree was reconstructed using 264 conserved proteins, 74 species, and 86,313 conserved amino acid positions, it was inferred with IQ-TREE under the PMSF approximation LG+F+R10+C20 model and ultrafast bootstrap as statistical support.

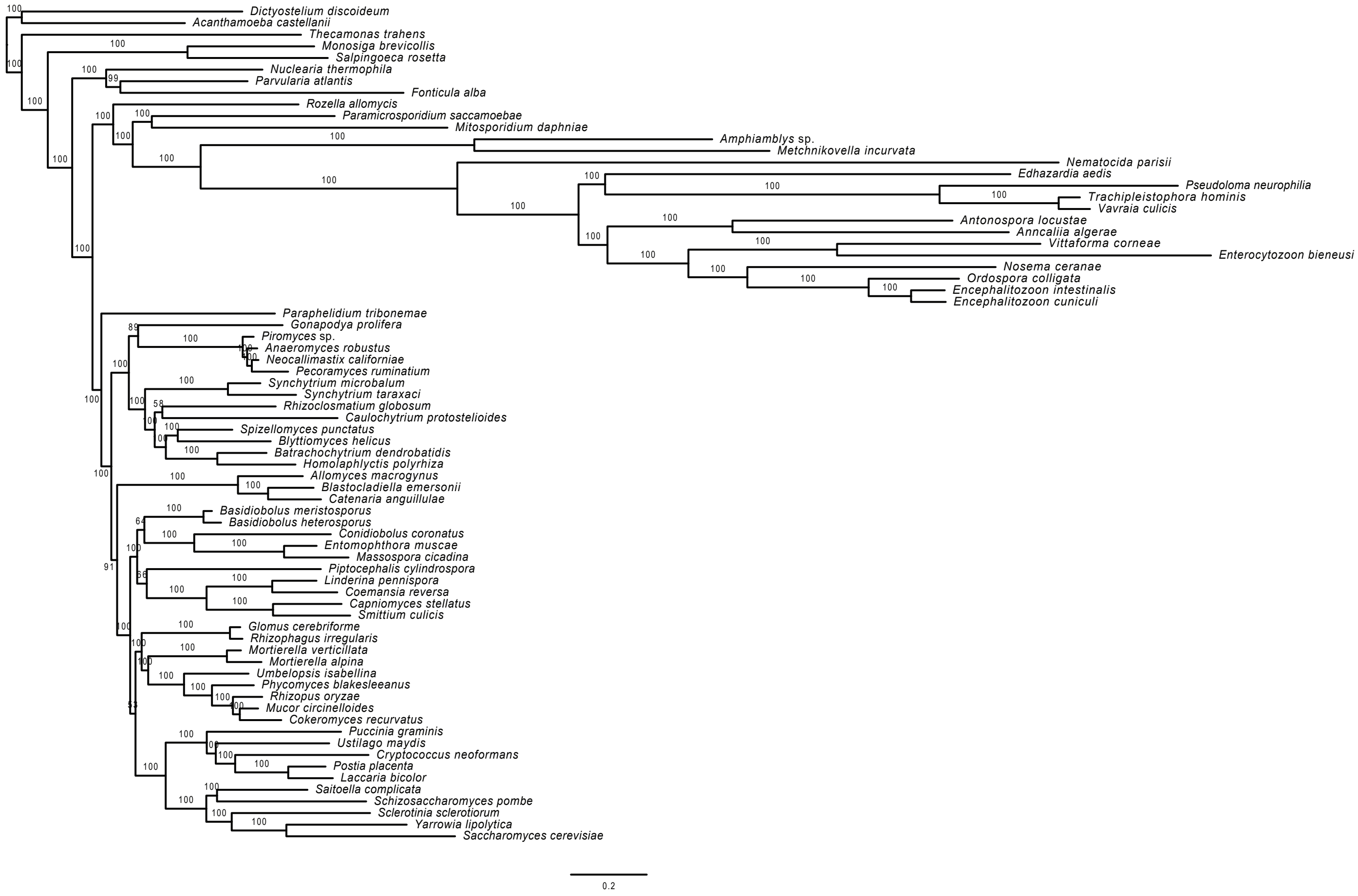


**Supplementary Fig. 2. e, Maximum likelihood tree based on the GBE protein dataset.** The tree was reconstructed using 264 conserved proteins, 72 species, and 84,949 conserved amino acid positions, it was inferred with IQ-TREE under the PMSF approximation LG+F+R9+C20 model and ultrafast bootstrap as statistical support.

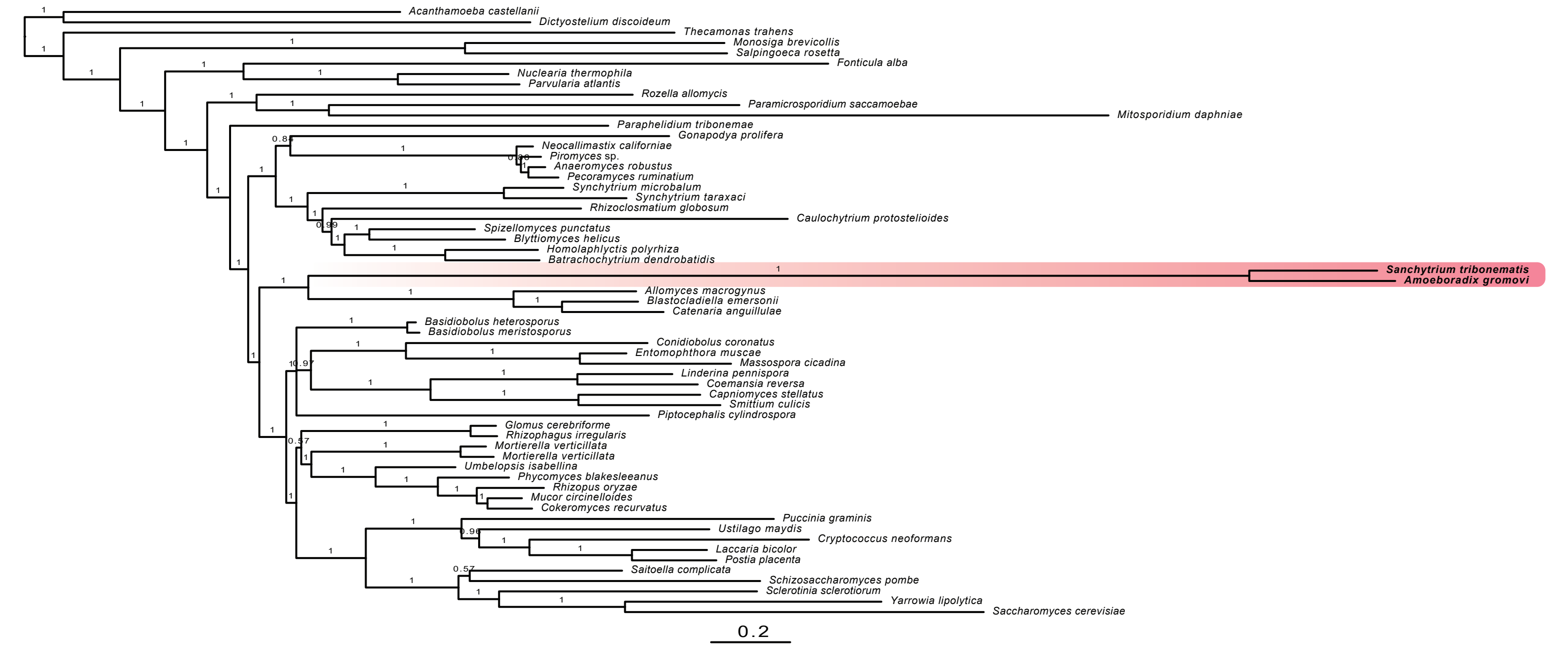


**Supplementary Fig. 2. f, Bayesian phylogenomic tree based on the BMC protein dataset.** The tree was reconstructed using 53 conserved proteins, 59 species, and 14,965 conserved amino acid positions, it was inferred using PhyloBayes under the CAT-GTR model with posterior probability as statistical support.

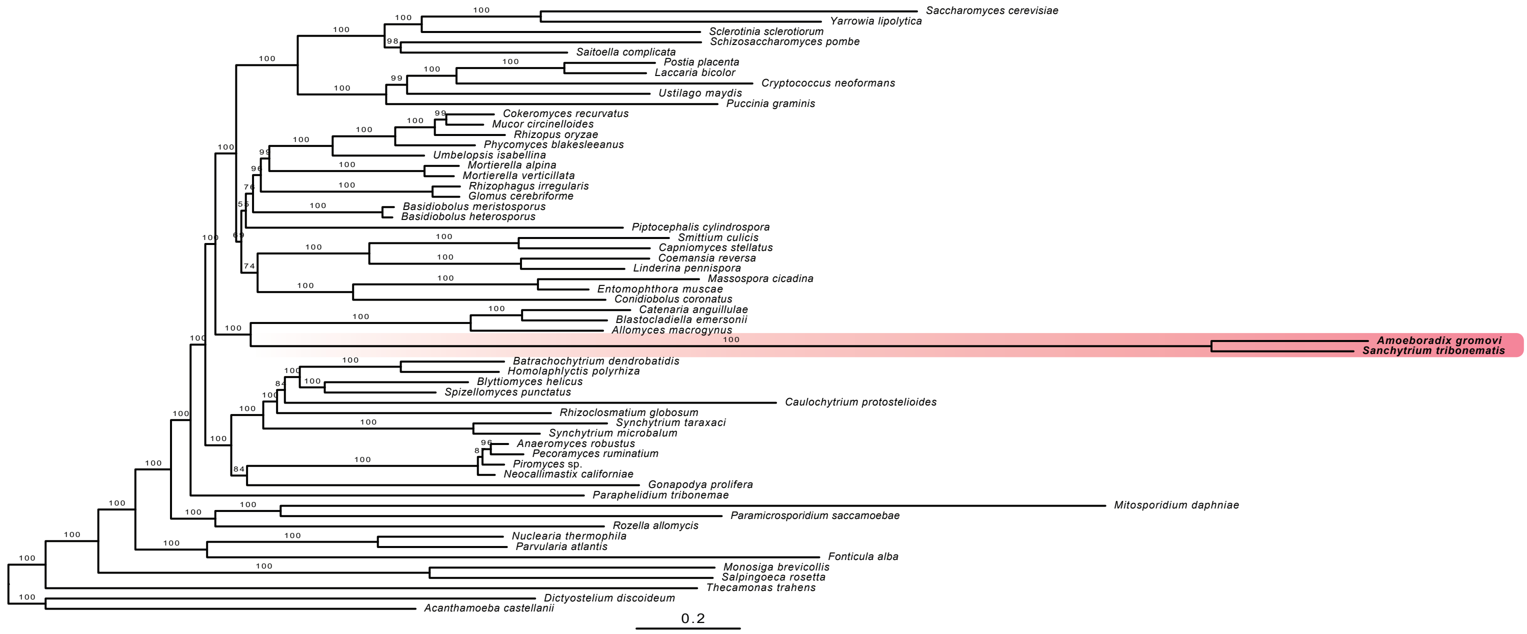


**Supplementary Fig. 2. g, Maximum likelihood tree based on the BMC protein dataset.** The tree was reconstructed using 53 conserved proteins, 59 species, and 14,965 conserved amino acid positions, it was inferred with IQ-TREE under the PMSF approximation of the LG+R7+C60 model and ultrafast bootstrap as statistical support.

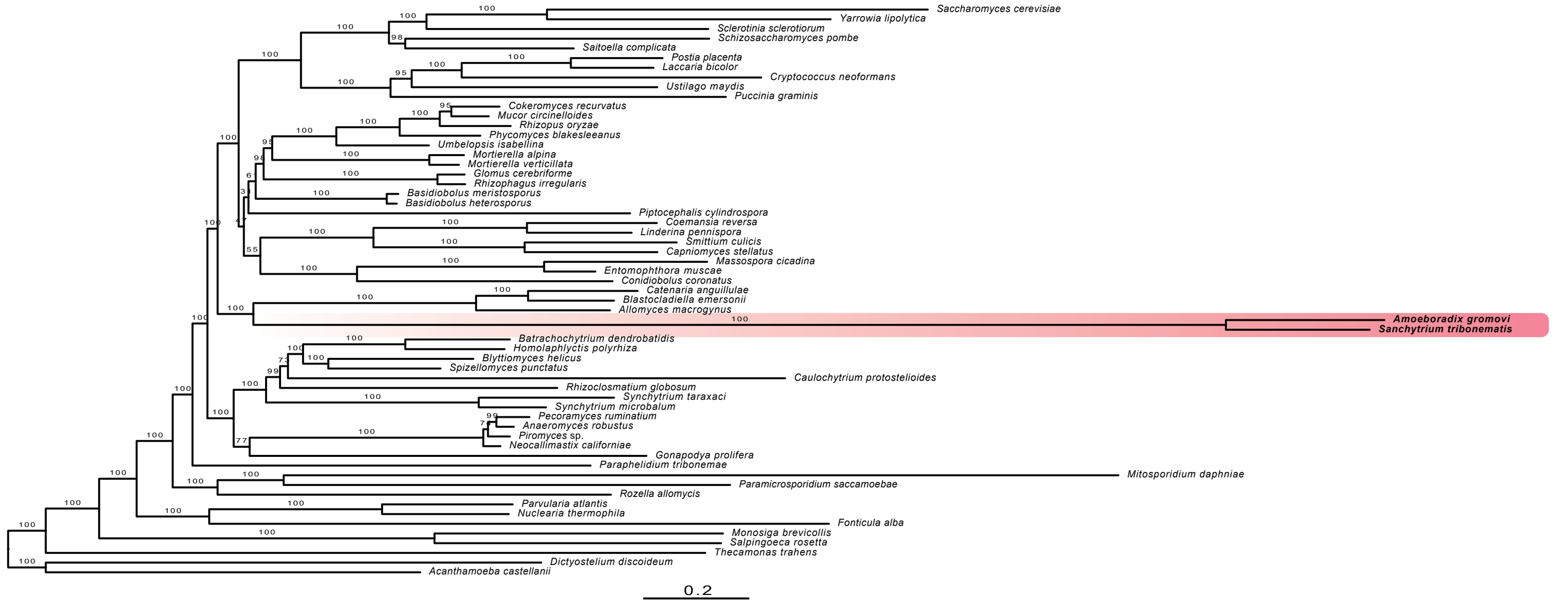


**Supplementary Fig. 2. h, Maximum likelihood tree based on the BMC protein dataset.** The tree was reconstructed using 53 conserved proteins, 59 species, and 14,965 conserved amino acid positions, it was inferred with IQ-TREE under the PMSF approximation of the LG+R7+C60 model and 100 conventional bootstrap replicates as statistical support.

| | *Cyberlindnera jadinii* | *Saccharomyces cerevisiae* | *Hanseniaspora gamundiae* | *Hanseniaspora singularis* | *Allomyces macrogynus* | ***Amoeboradix gromovi*** | ***Sanchytrium tribonematis*** |
|---|---|---|---|---|---|---|---|
| Sen15 | Present | Present | Present | Absent | Absent | Absent | Absent |
| Mcm21 | Present | Present | Present | Absent | Absent | Absent | Absent |
| Sld2 | Present | Present | Present | Absent | Absent | Absent | Absent |
| Cdc13 | Present | Present | Present | Absent | Absent | Absent | Absent |
| Mcm22 | Absent | Present | Present | Absent | Absent | Absent | Absent |
| Abf1 | Absent | Present | Present | Absent | Absent | Absent | Absent |
| Rfa3 | Absent | Present | Present | Absent | Absent | Absent | Absent |
| Csm2 | Absent | Present | Present | Absent | Absent | Absent | Absent |
| Fyv6 | Present | Present | Absent | Absent | Absent | Absent | Absent |
| Rif1 | Present | Present | Present | Absent | Absent | Absent | Absent |
| Pds1 | Present | Present | Present | Absent | Absent | Absent | Absent |
| Mms22 | Present | Present | Present | Absent | Absent | Absent | Absent |
| Eco1 | Present | Present | Present | Absent | Present | Absent | Absent |
| Mms4 | Present | Present | Present | Absent | Absent | Absent | Absent |
| Mrc1 | Present | Present | Present | Absent | Absent | Absent | Absent |
| Nse1 | Present | Present | Present | Absent | Absent | Absent | Absent |
| Nse3 | Present | Present | Present | Absent | Absent | Absent | Absent |
| Nse5 | Present | Present | Present | Absent | Absent | Absent | Absent |
| Nup120 | Present | Present | Present | Absent | Absent | Absent | Absent |
| Nup133 | Present | Present | Present | Absent | Absent | Absent | Absent |
| Nup60 | Present | Present | Present | Absent | Absent | Absent | Absent |
| Rad9 | Present | Present | Present | Absent | Absent | Absent | Absent |
| Slx4 | Present | Present | Present | Absent | Absent | Absent | Absent |
| Snf6 | Present | Present | Present | Absent | Absent | Absent | Absent |
| Tah11 | Present | Present | Present | Absent | Absent | Absent | Absent |
| Eaf6 | Absent | Present | Present | Absent | Present | Absent | Absent |
| Pol32 | Present | Present | Present | Absent | Absent | Absent | Absent |
| Def1 | Present | Present | Present | Absent | Absent | Absent | Absent |
| Tdp1 | Present | Present | Present | Absent | Absent | Present | Absent |
| Pol4 | Present | Present | Present | Absent | Present | Present | Present |
| Psy3 | Present | Present | Absent | Absent | Absent | Absent | Absent |
| Hpr1 | Present | Present | Absent | Absent | Absent | Present | Present |
| Sae3 | Present | Present | Absent | Absent | Absent | Absent | Absent |
| Kre29 | Absent | Present | Absent | Absent | Absent | Absent | Absent |
| Eaf5 | Absent | Present | Absent | Absent | Absent | Absent | Absent |
| Mms1 | Absent | Present | Absent | Absent | Absent | Absent | Absent |
| Nej1 | Absent | Present | Absent | Absent | Absent | Absent | Absent |
| Sir4 | Absent | Present | Absent | Absent | Absent | Absent | Absent |
| Lif1 | Absent | Present | Absent | Absent | Absent | Absent | Absent |
| Xrs2 | Absent | Present | Absent | Absent | Absent | Absent | Absent |
| Eaf7 | Absent | Present | Absent | Absent | Present | Absent | Absent |
| Mag1 | Present | Present | Absent | Absent | Absent | Absent | Absent |
| Eaf3 | Present | Present | Absent | Absent | Present | Present | Present |
| Pcd1 | Present | Present | Absent | Absent | Present | Present | Present |
| Phr1 | Present | Present | Absent | Absent | Present | Present | Present |
| Mgt1 | Absent | Present | Absent | Present | Present | Present | Present |
| Lrs4 | Absent | Present | Present | Absent | Absent | Absent | Absent |

Legend: Present (green); Absent (white)

**Supplementary Fig. 3.** Genome maintenance and DNA repair genes in *Amoeboradix* and *Sanchytrium*, compared with a close relative (the blastoclad *Allomyces*) and several slow- and fast-evolving yeasts. Based on the study by Steenwyk et al. (2019) Extensive loss of cell-cycle and DNA repair genes in an ancient lineage of bipolar budding yeasts. PLoS Biol 17(5): e3000255.

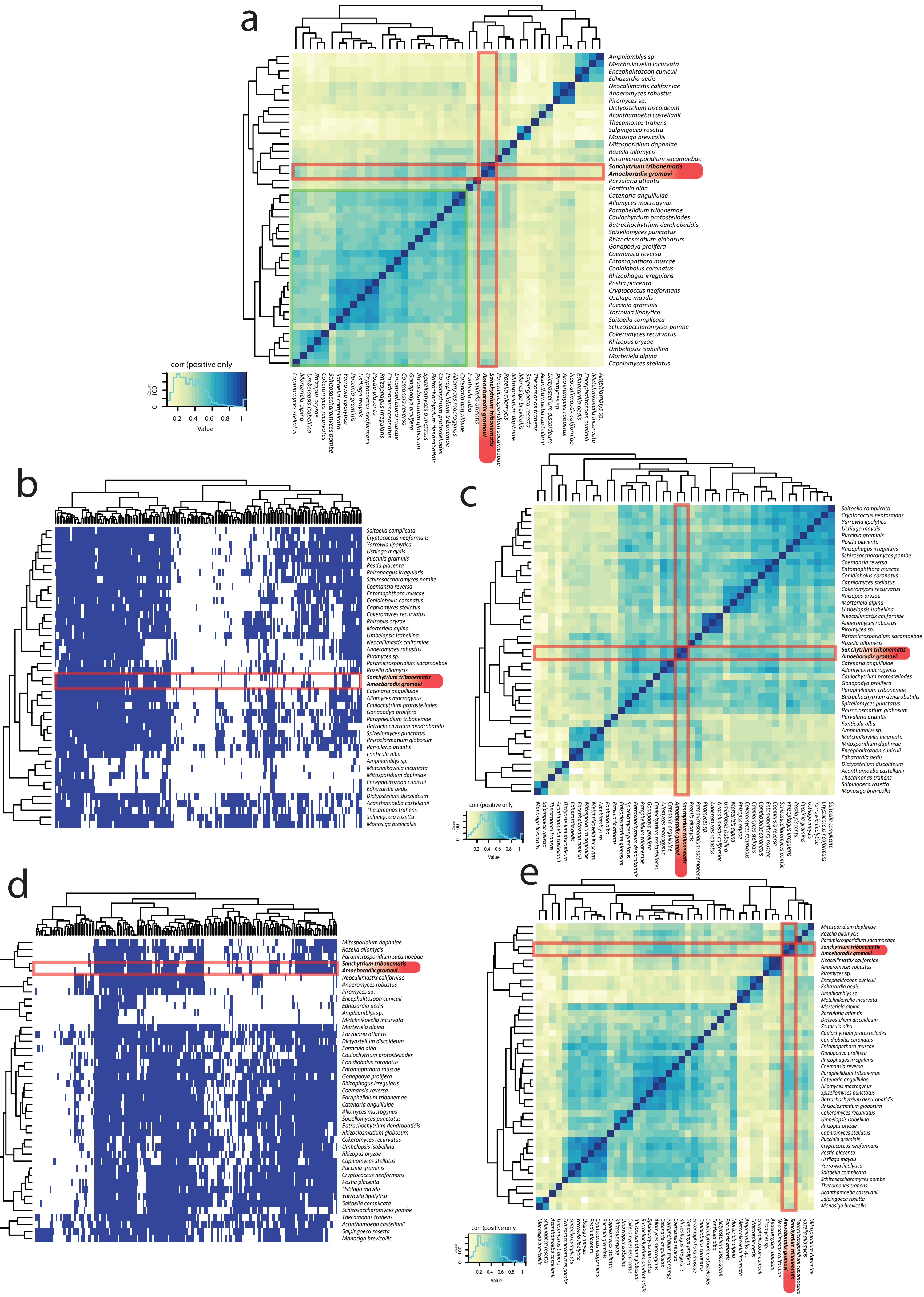


**Supplementary Fig. 4. Comparative analyses of primary metabolism. a,** Clustering of 43 eukaryotic species according to their profile of presence/absence of the same 8 primary metabolism categories for a total of 1158 COGs. Color code is scaled to display positive Pearson's r (0 to 1). (**b** and **d**) Binary heat maps of orthologs (COGs) in 43 eukaryotic genomes/transcriptomes. (**c** and **e**) Clustering of 43 eukaryotic genomes/transcriptomes according to their profile of presence/absence color coded to display positive Pearson's r (0 to 1) (**b** and **c**) Carbohydrate transport and metabolism (205 OGs, category G). (**d** and **e**) Lipid transport and metabolism (201 OGs, category I).

a

b

c

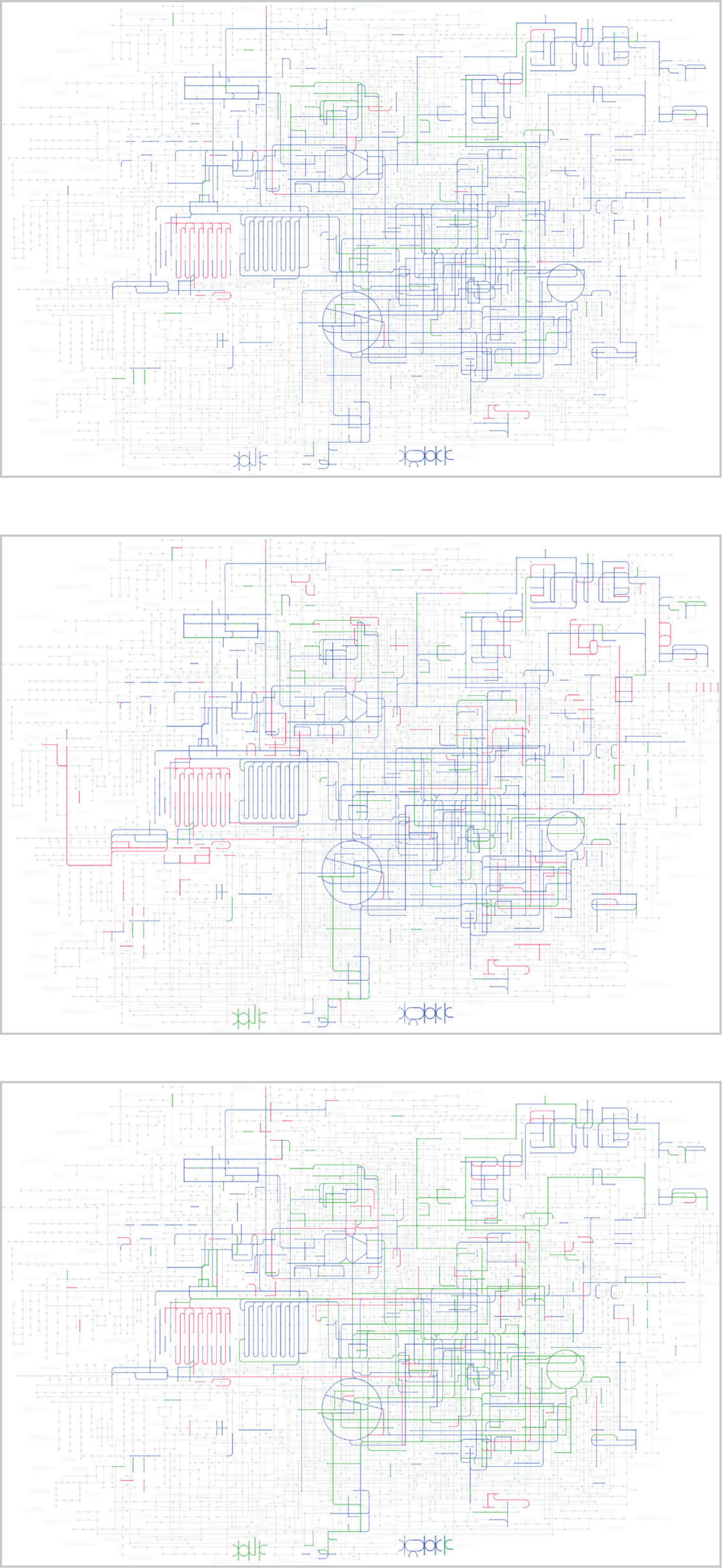

**Supplemetary Fig. 5. a - c, KEGG Pathways metabolic comparison.** (**a** - **c**) KEGG metabolic pathways map01100. **a**, *Sanchytrium tribonematis* (green) vs *Amoeboradix gromovi* (pink); both (blue). **b**, *Sanchytrium tribonematis* (green) vs *Allomyces macrogynus* (pink); both (blue). **c**, *Sanchytrium tribonematis* (green) vs *Rozella allomycis* (pink); both (blue).

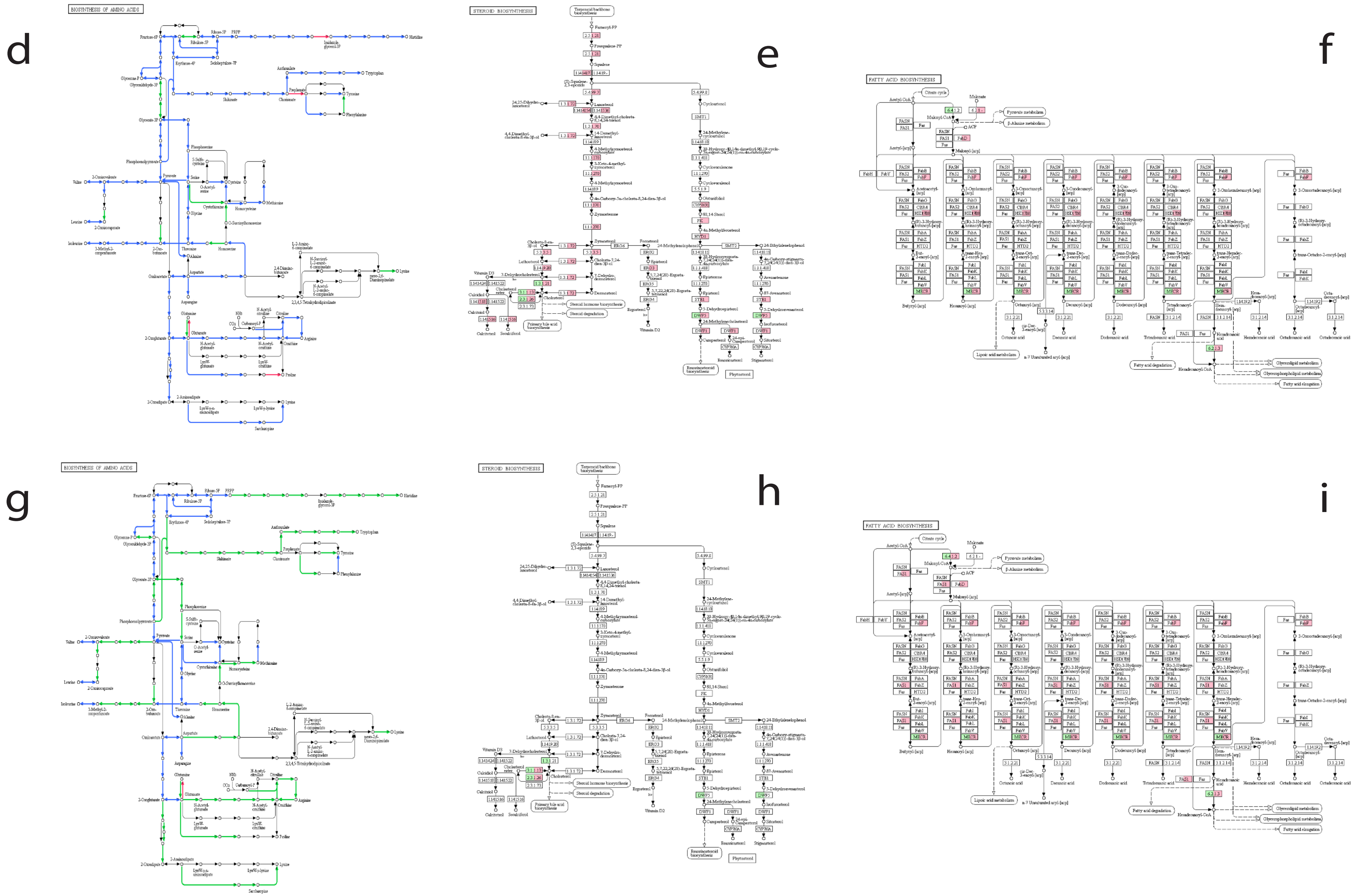


**Supplementary Fig. 5. d -i, KEGG Pathways metabolic comparison.** (**d** - **f**) KEEG specific metabolic pathways of *Sanchytrium tribonematis* (green) vs *Allomyces macrogynus* (pink); both (blue). (**g** - **i**) KEEG specific metabolic pathways of *Sanchytrium tribonematis* (green) vs *Rozella allomycis* (pink); both (blue). (**d** and **g**) KEGG Amino acid synthesis map01230. (**e** and **h**) KEGG Steroid biosynthesis map00100. (**f** and **i**) KEGG Fa  y acid biosynthesis map00061.

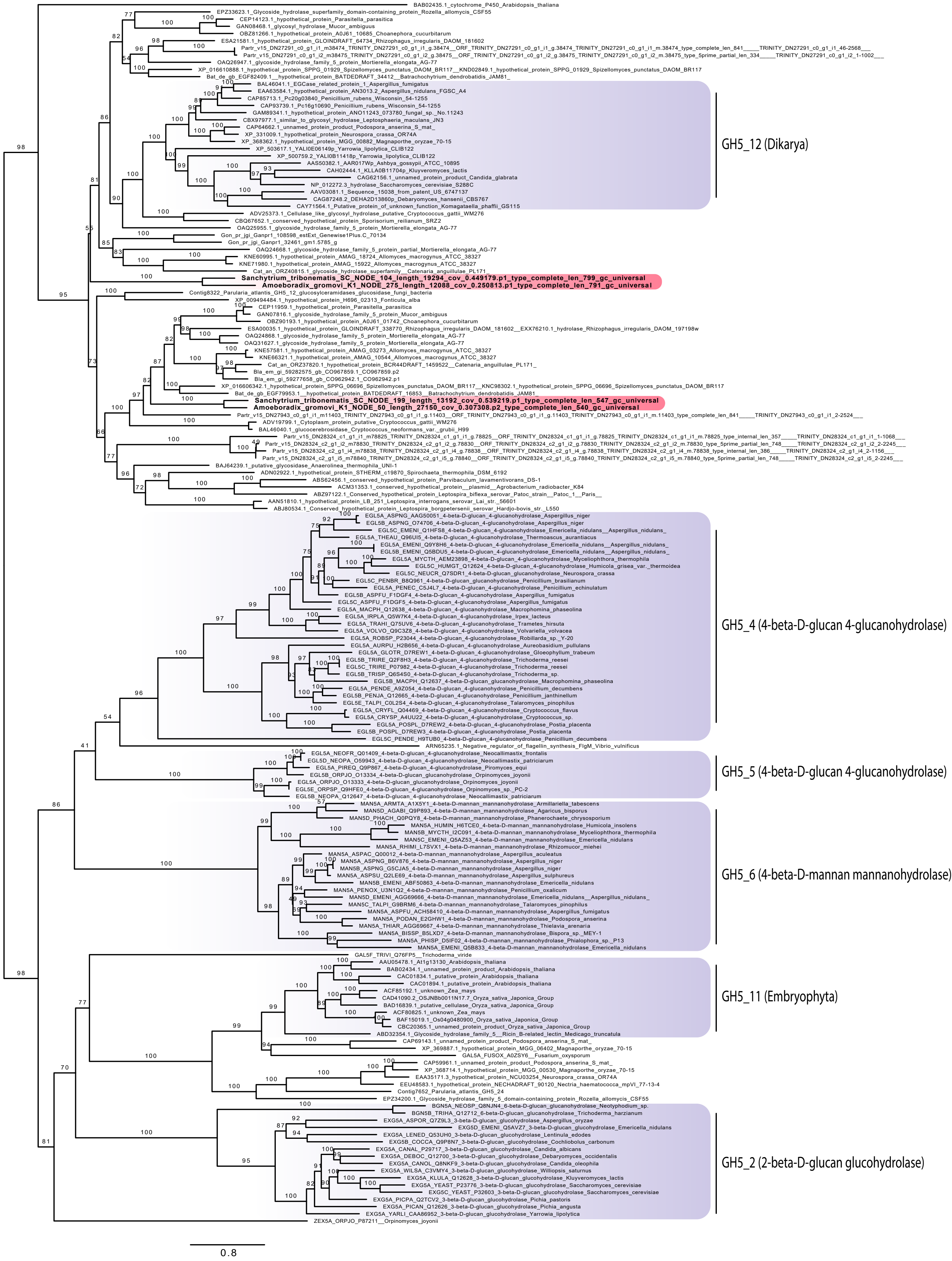


**Supplementary Fig. 6. Maximum likelihood tree of the GH5 cellulase family.** The tree was reconstructed based on the dataset of Torruella et al. 2018, 170 sequences and 4,721 amino acid positions with the LG+R7 model for ML. Ultrafast bootstrap (ufbs) support values are indicated over the branches.

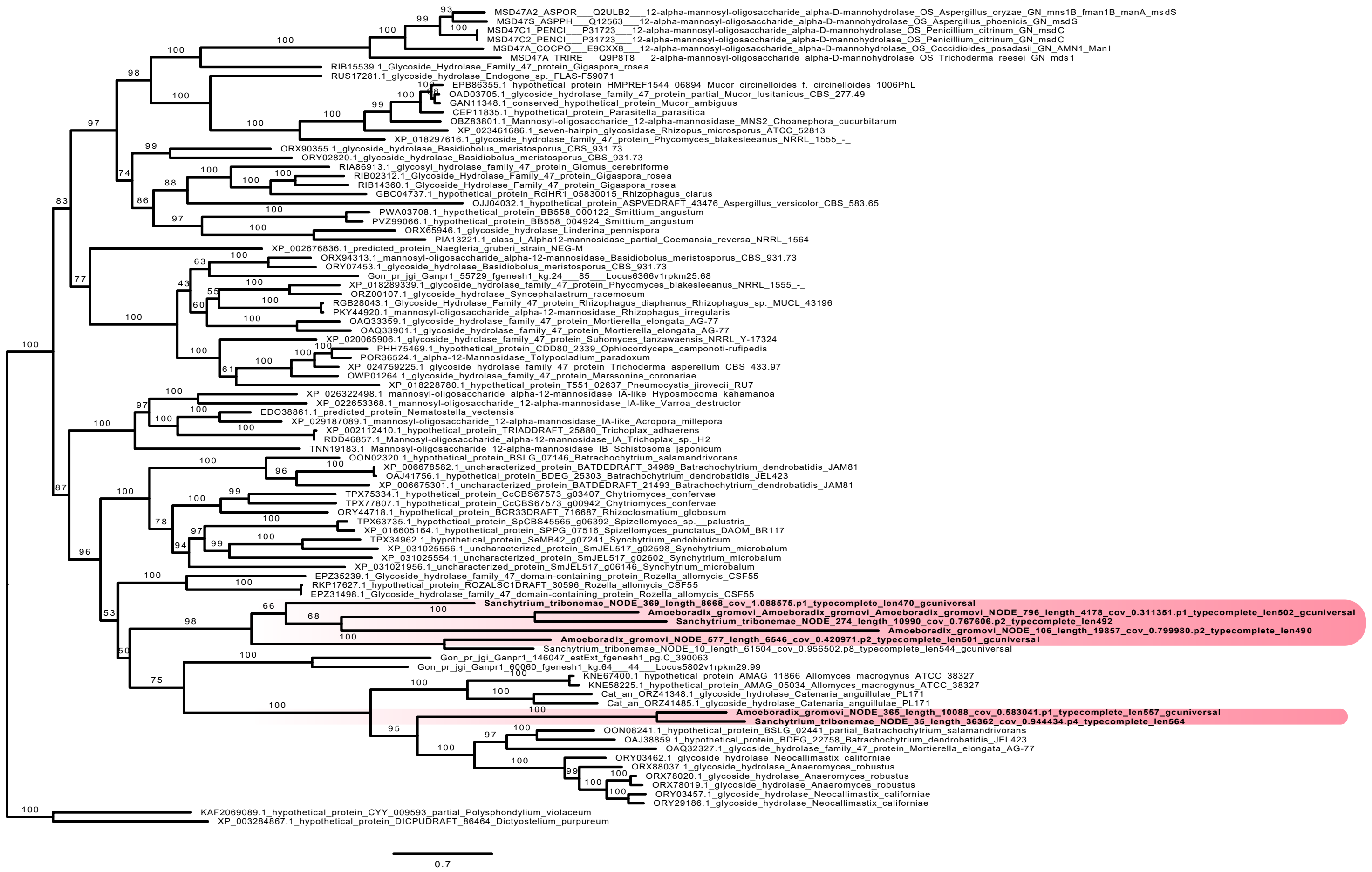


**Supplementary Fig. 7.** Maximum likelihood tree of the GH47 alpha-1,2-mannosidase family of hemicellulose degradation, containing a total of 90 sequences and 1,910 amino acid positions with the LG+F+R6 model for ML. Ultrafast bootstrap (ufbs) support values are indicated over the branches.

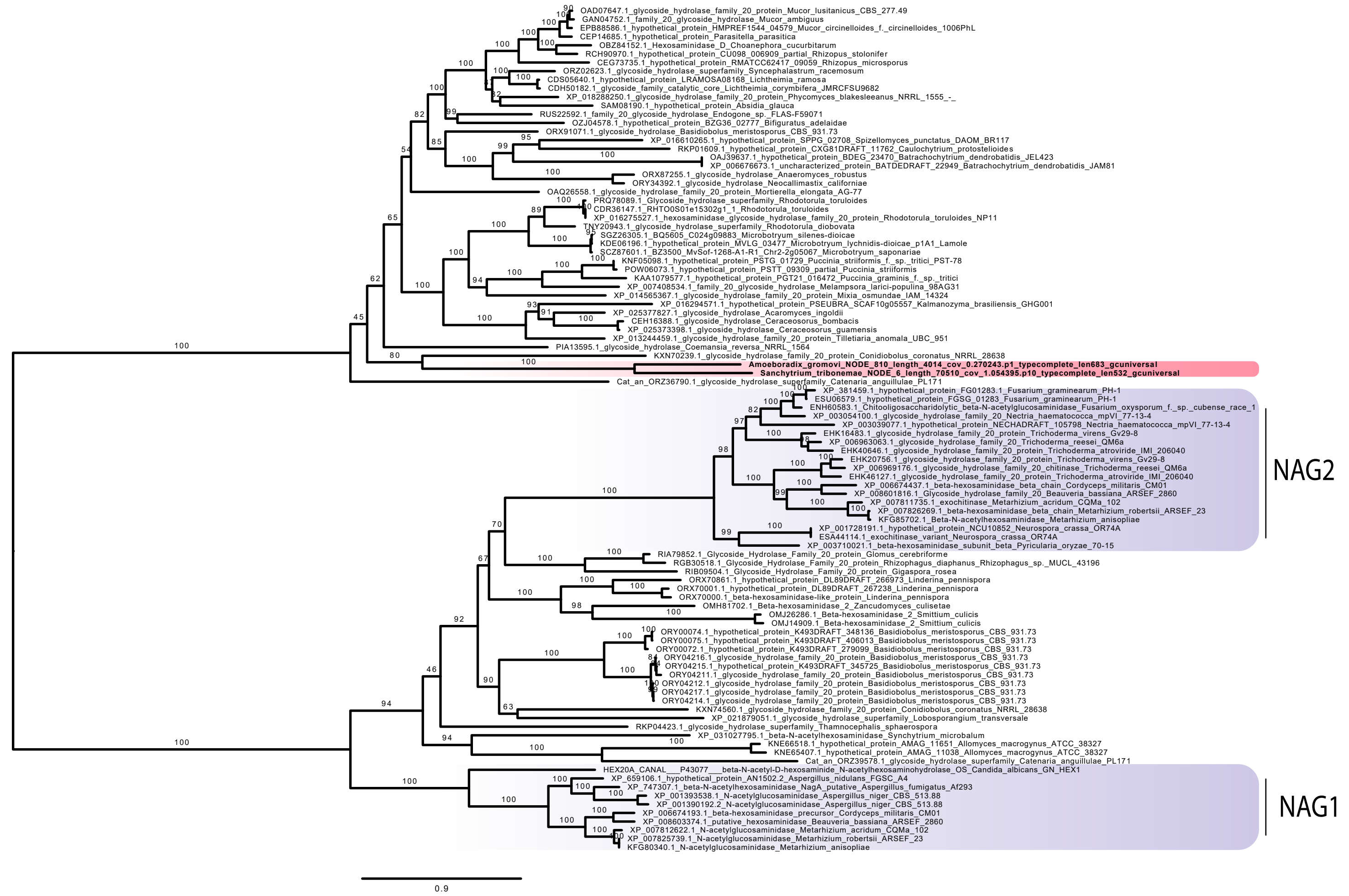


**Supplementary Fig. 8.** Maximum likelihood tree of family GH20 b-N-acetylhexosaminidases (NAGase) containing a total of 98 sequences and 2,031 amino acid positions with the LG+F+R7 model for ML. The tree was reconstructed based on the dataset of de Oliveira et al. 2018. Ultrafast bootstrap (ufbs) support values are indicated over the branches.

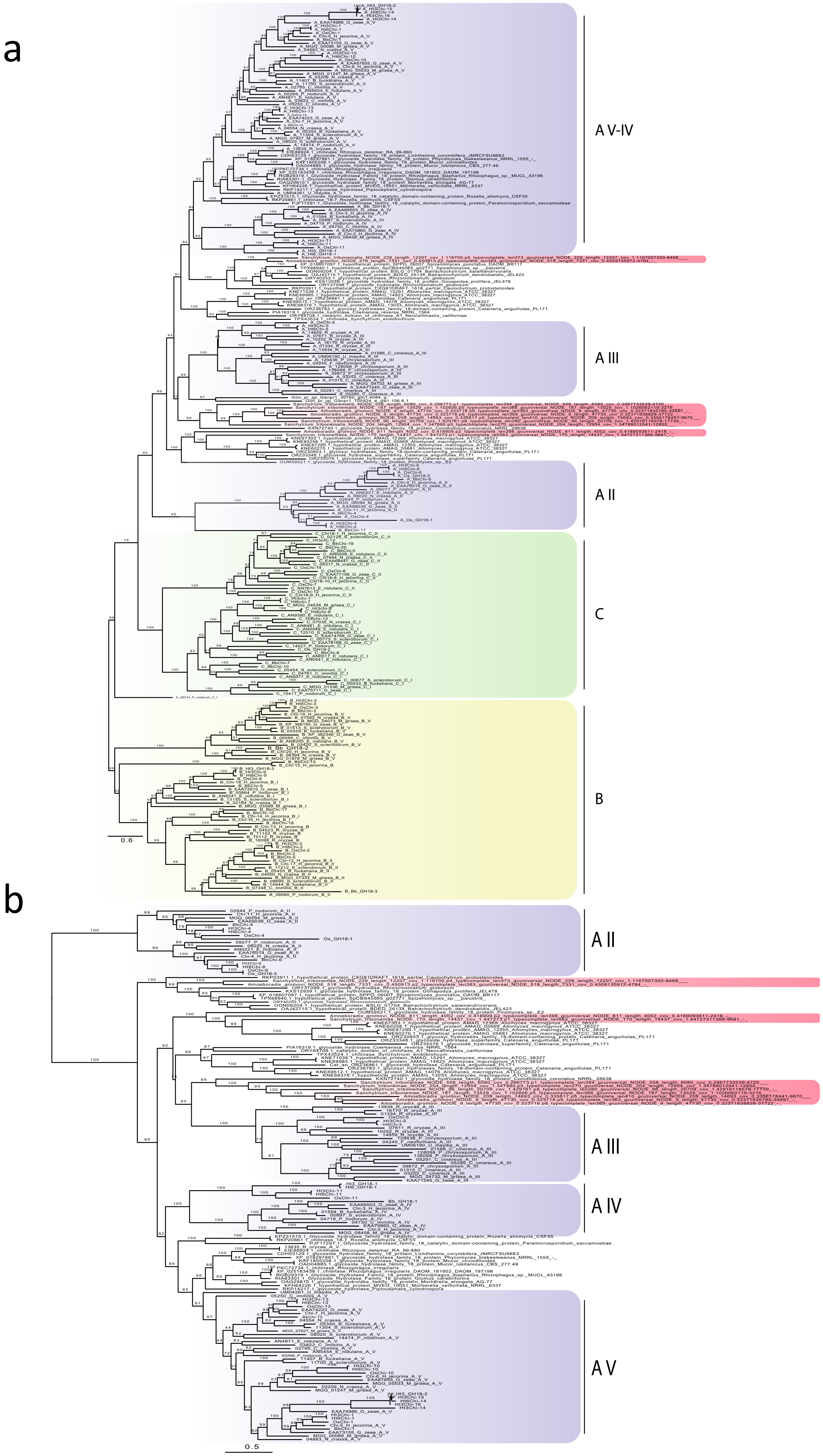


**Supplementary Fig. 9. Maximum likelihood trees of the GH18 chytinase family.** The trees were reconstructed based on the dataset of Agrawal et al. 2015. (A) Maximum likelihood tree of 262 sequences and 5,761 amino acid positions with the LG+R8 model for ML. (B) Maximum likelihood tree of family GH18A of 152 sequences and 2,746 amino acid positions with the LG+R6 model for ML. Ultrafast bootstrap (ufbs) support values are indicated over the branches.

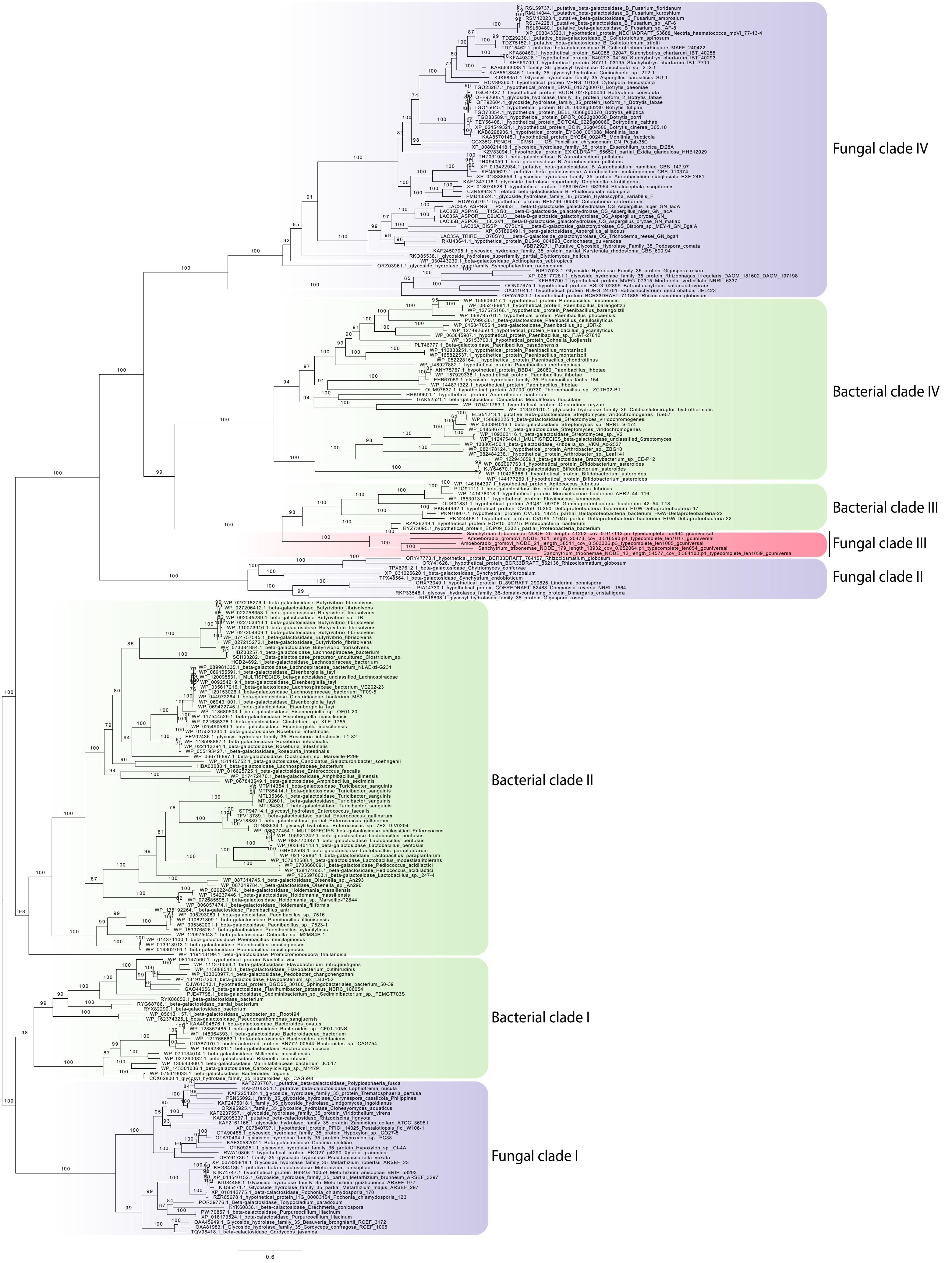


**Supplementary Fig. 10. Maximum likelihood tree of the GH35 β-galactosidase family.** Maximum likelihood tree of 248 sequences and 2,424 amino acid positions with the LG+R7 model for ML. Ultrafast bootstrap (ufbs) support values are indicated over the branches.

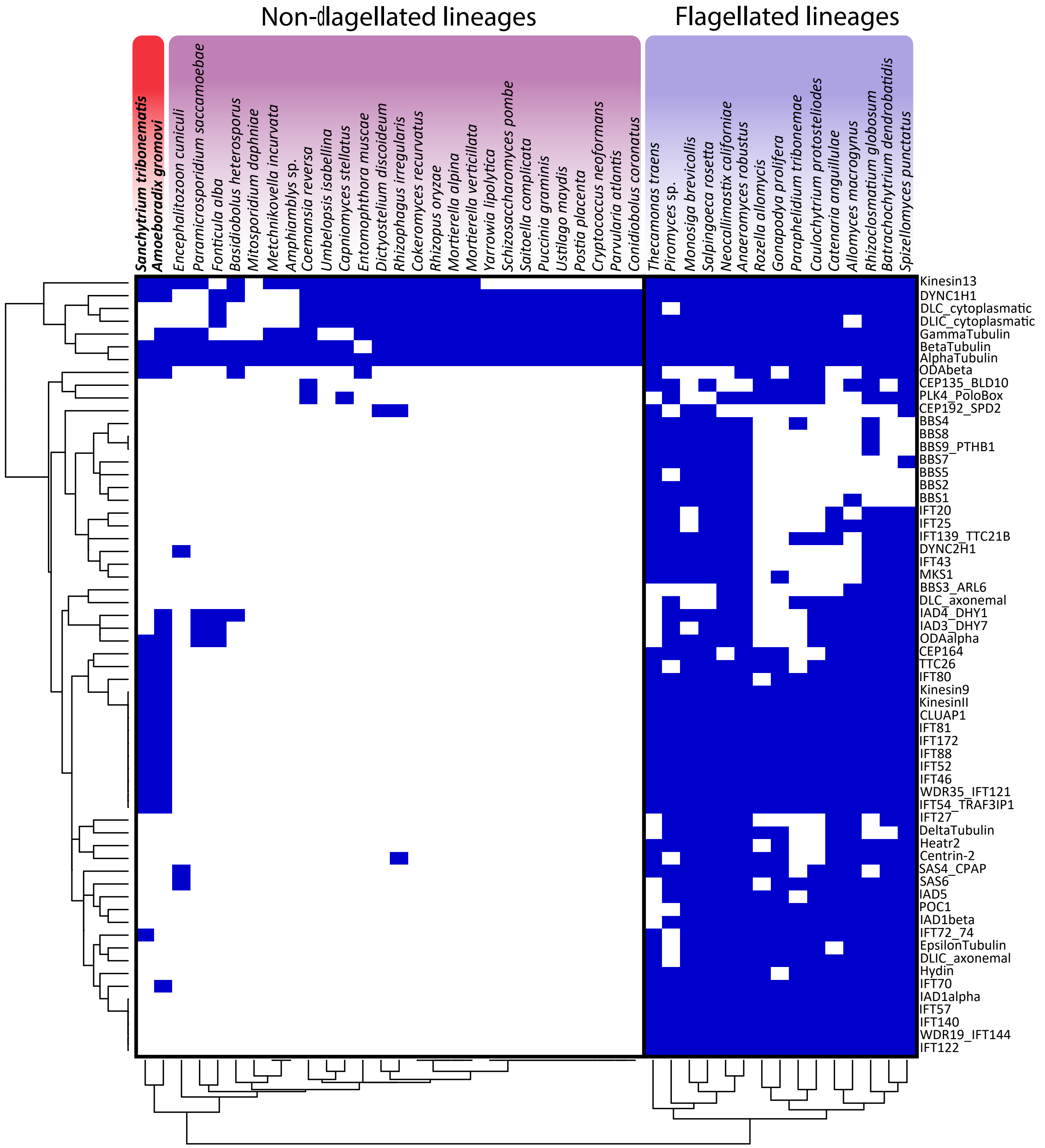


**Supplementary Fig. 11. Flagellum toolkit reduction.** Presence/absence heatmaps of 61 flagellum-specific proteins in 43 eukaryotic flagellated (purple) and non-flagellated (pink) lineages. Heatmap clustered by similarity showing sanchytrids in an intermediate position between flagellated and non-flagellated lineages. Gene presence is depicted in blue and absence is depicted in white.

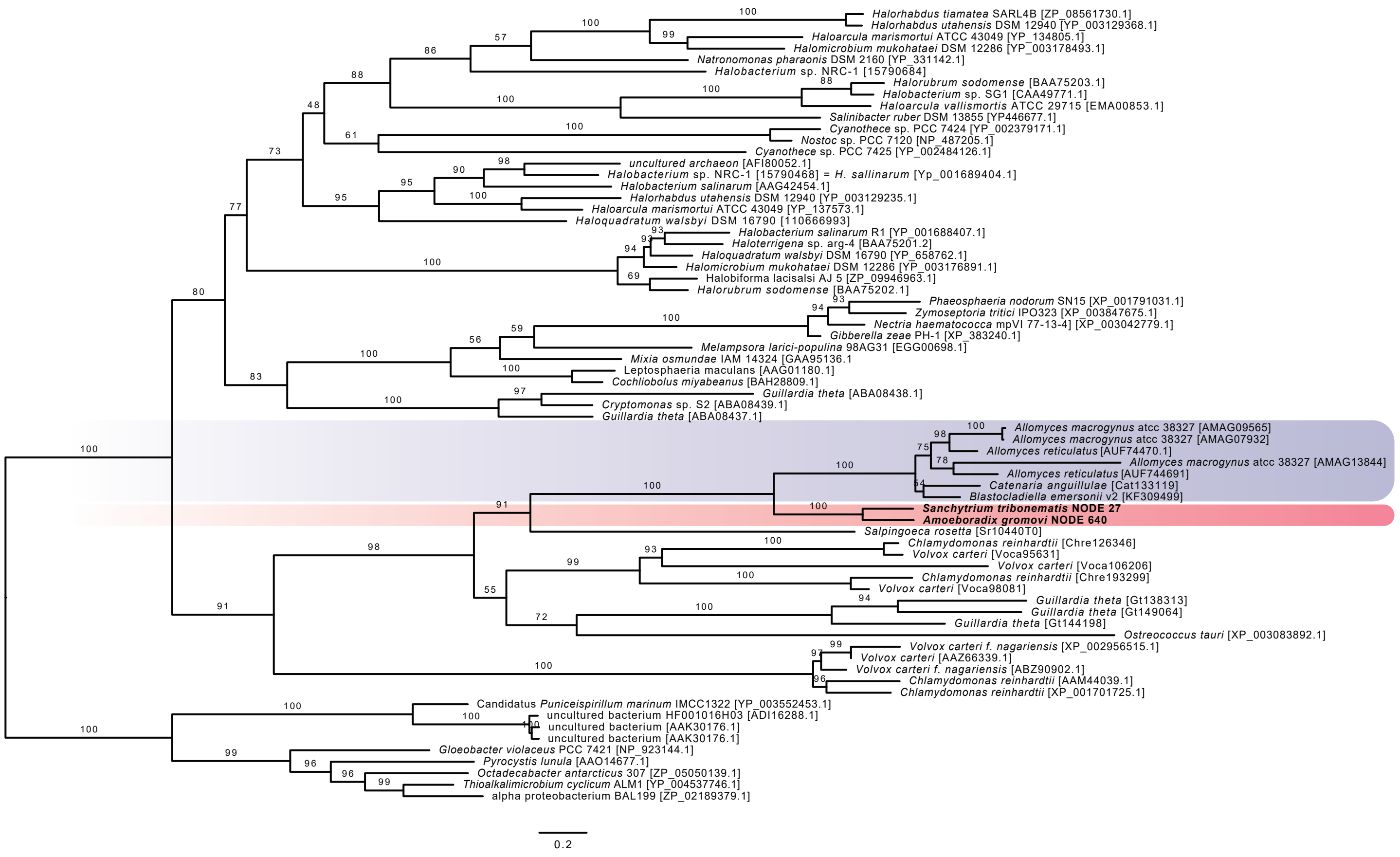


**Supplementary Fig. 12. a, Maximum likelihood tree based on the protein dataset of Avelar et al. (2014).** Reconstruction of the Type I rhodopsin domain of the BeGC1 gene-fusion, with 69 sequences and and 416 amino acid positions, it was inferred with IQ-TREE under the LG+F+I+G4 model and ultrafast bootstrap as statistical support.

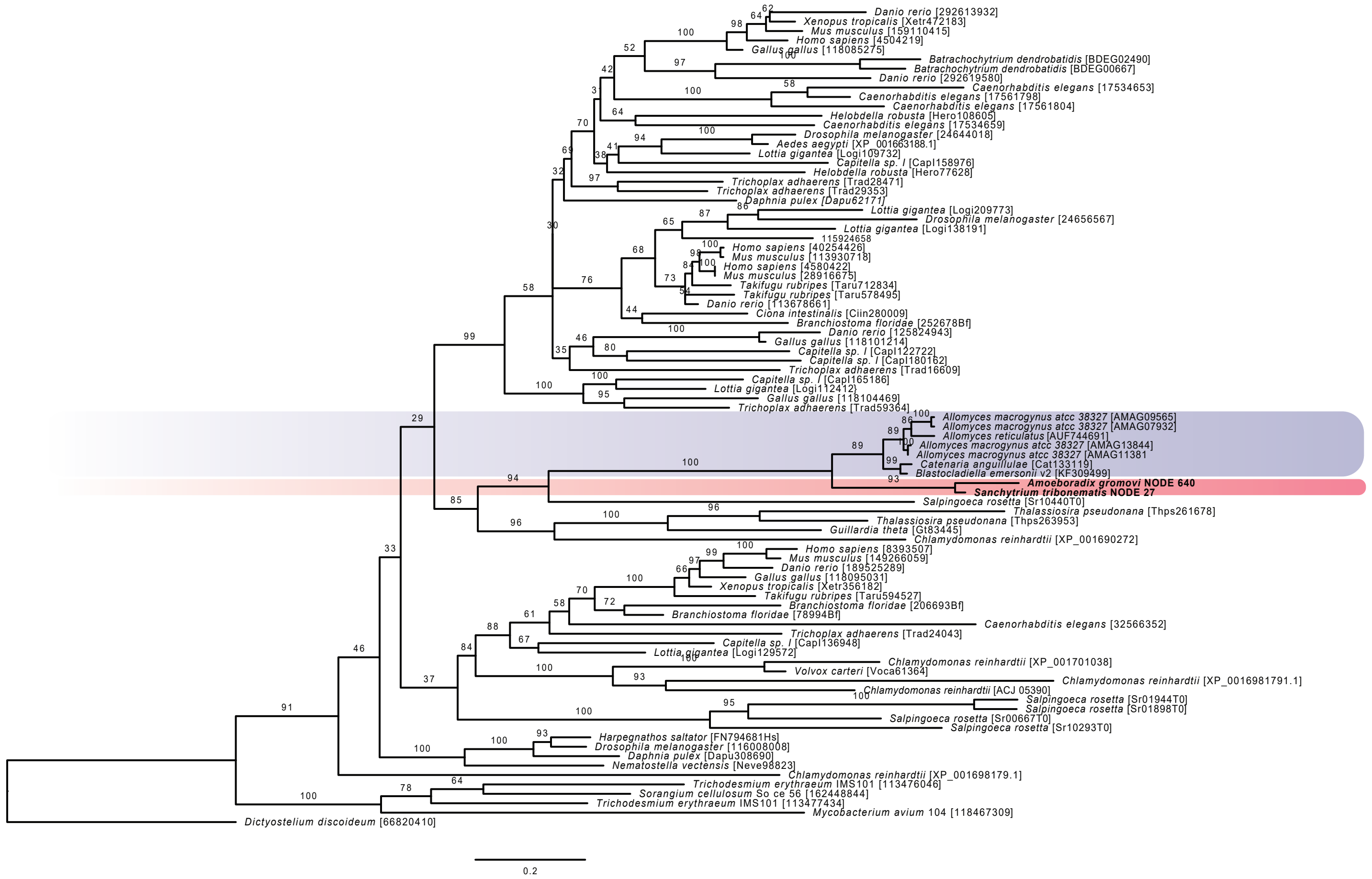


**Supplementary Fig. 12. b, Maximum likelihood tree based on the protein dataset of Avelar et al. (2014).** Reconstruction of the GC1 guanylyl-cyclase domain of the BeGC1 gene-fusion, with 87 sequences and 180 amino acid positions, it was inferred with IQ-TREE under the LG+G4 model and ultrafast bootstrap as statistical support.

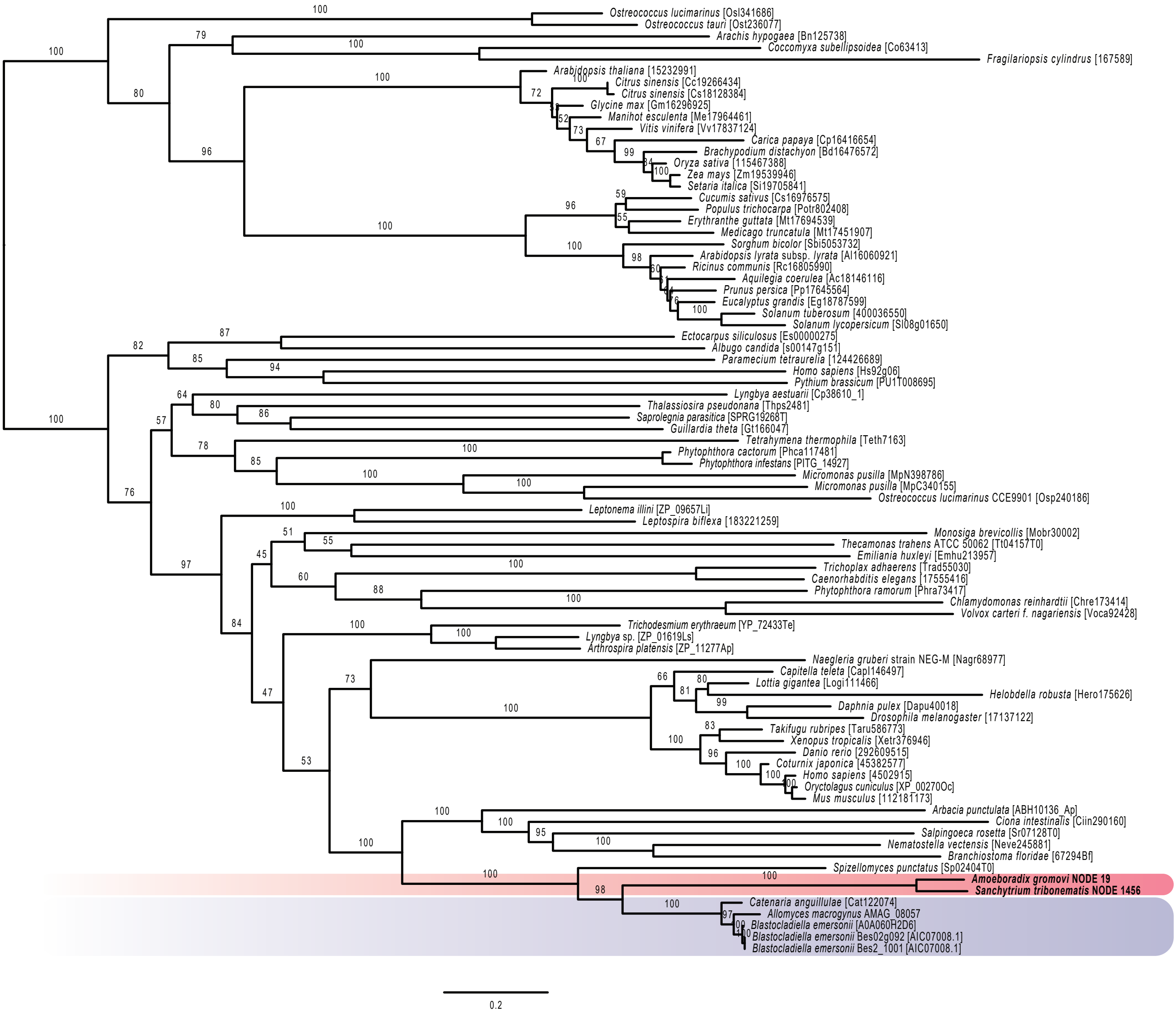


**Supplementary Fig. 12. c, Maximum likelihood tree based on the protein dataset of Avelar et al. (2014).** Reconstruction of the cyclic nucleotide gated channel BeCNG1, and 301 amino acid positions, it was inferred with IQ-TREE under the LG+F+I+G4 model and ultrafast bootstrap as statistical support.

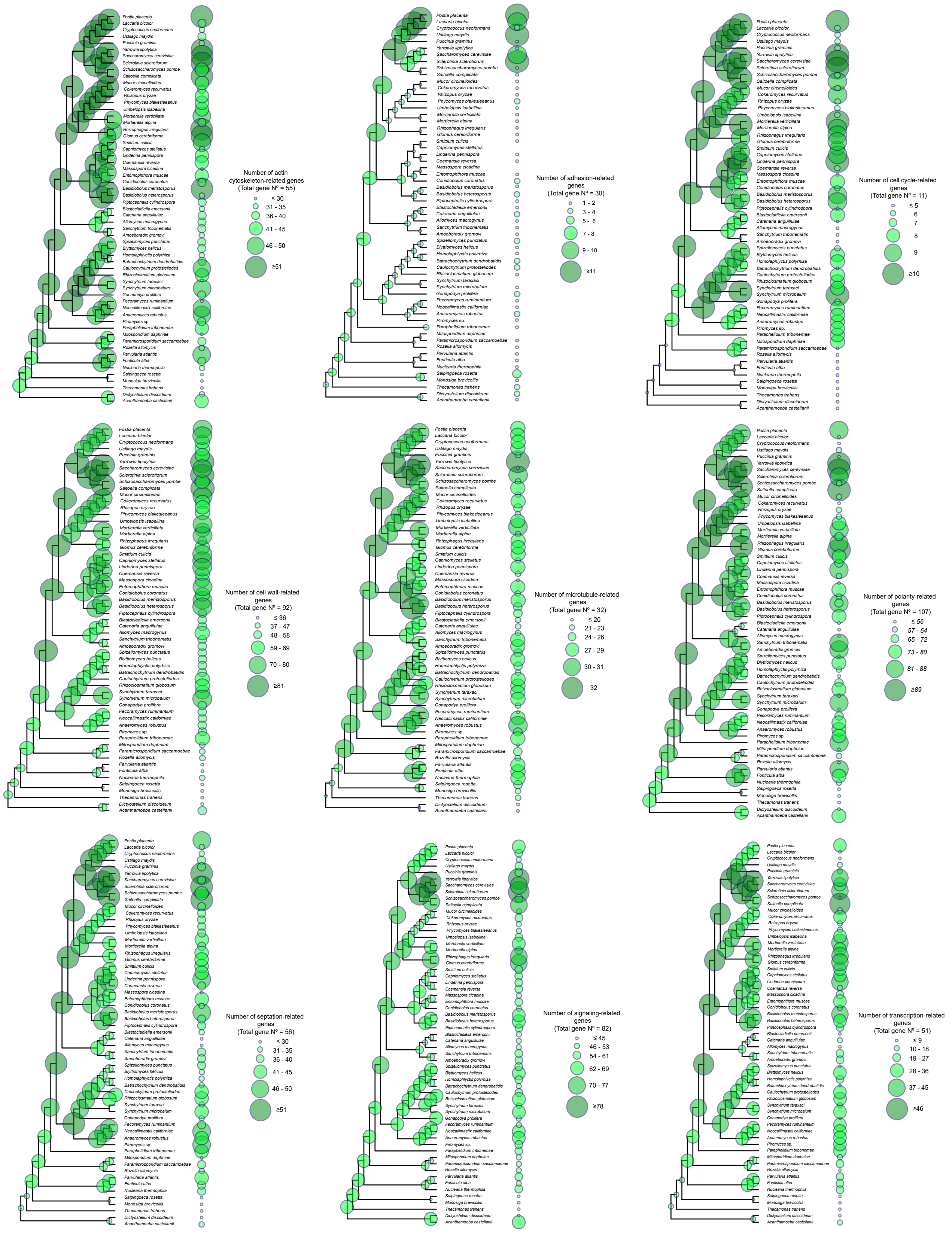

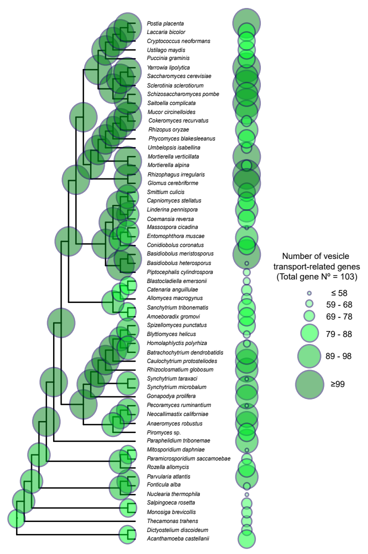


**Supplementary Fig. 13. Evolution of hyphal multicellularity in Holomycota.** Cladograms of Holomycota according to the GBE59 ML phylogenomic reconstruction for 10 different hyphal morphogenesis functional categories. Bubble size on the nodes and tips are represent the total number of reconstructed ancestral and extant hyphal multicellularity related gene proteins for each functional category.

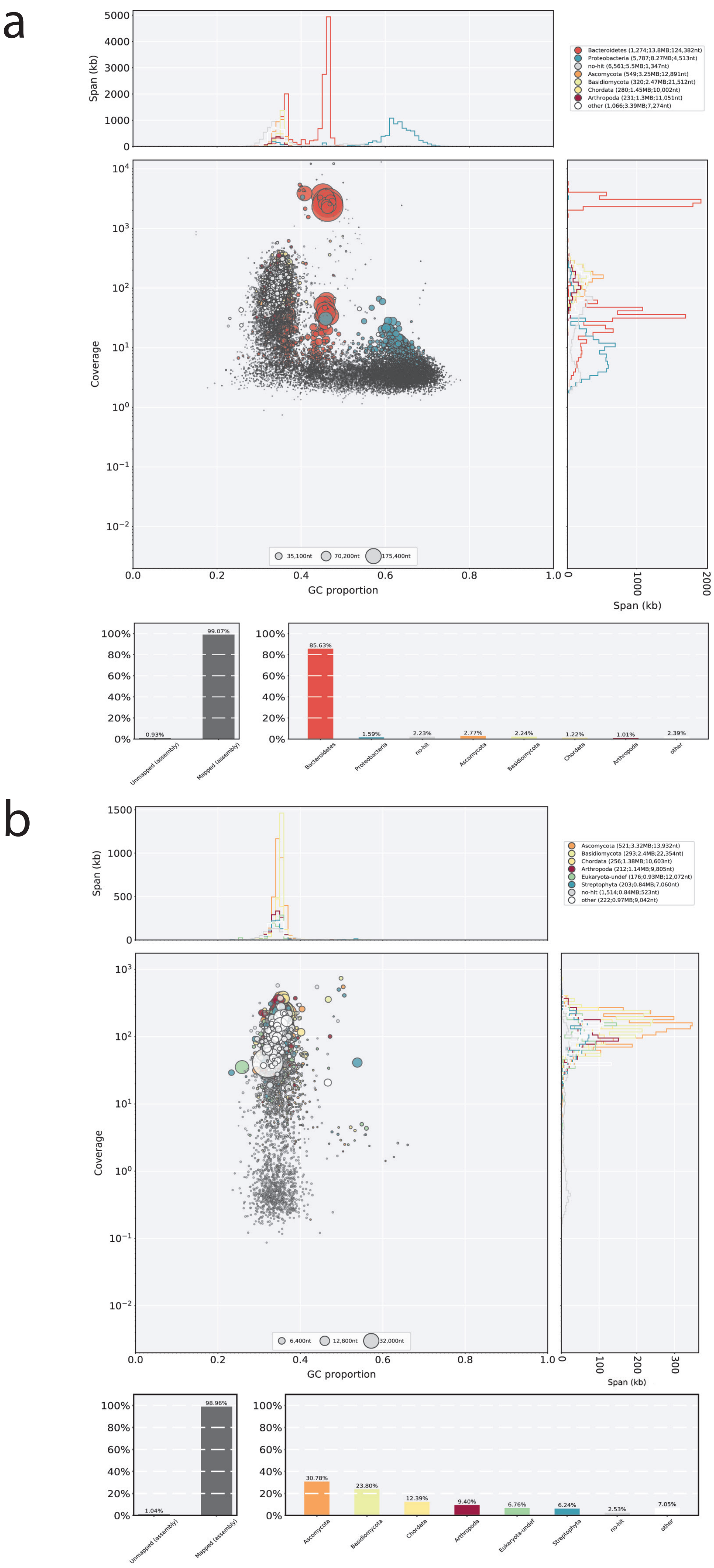


**Supplementary Fig. 14. a-b,** Blobplots showing the results before (**a**) and after (**b**) bacterial decontamination of the genome of *Sanchytrium tribonematis*. Taxon-annotated scatter plots decorated with coverage and GC content, the legend represents the taxonomic affiliation of sequences (phylum level) and lists count, total span and N50 by taxonomic group. Histograms show the proportion of reads of a library that are unmapped or mapped, showing the percentage of mapped reads by phylum.

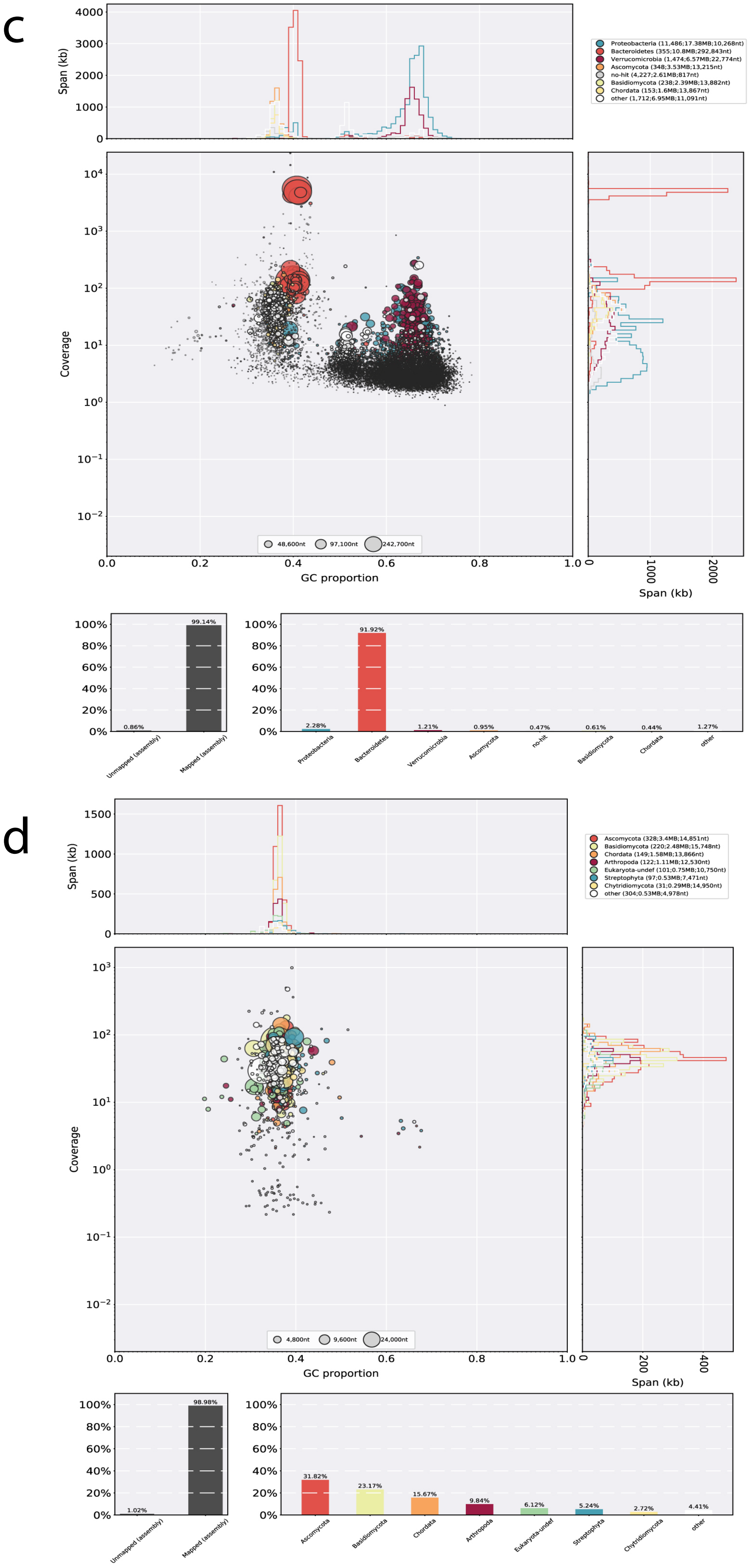


**Supplementary Fig. 14. c-d,** Blobplots showing the results before (**c**) and after (**d**) bacterial decontamination of the genome of *Amoeboradix gromovi*. Taxon-annotated scatter plots decorated with coverage and GC content, the legend represents the taxonomic affiliation of sequences (phylum level) and lists count, total span and N50 by taxonomic group. Histograms show the proportion of reads of a library that are unmapped or mapped, showing the percentage of mapped reads by phylum.
